# *Methanomethylophilus alvus Mx1201* provides basis for mutual orthogonal pyrrolysyl tRNA/aminoacyl-tRNA synthetase pairs in mammalian cells

**DOI:** 10.1101/371757

**Authors:** Birthe Meineke, Johannes Heimgärtner, Lorenzo Lafranchi, Simon J Elsässer

## Abstract

Genetic code expansion via stop codon suppression is a powerful technique for engineering proteins in mammalian cells with site-specifically encoded non-canonical amino acids (ncAAs). Current methods rely on very few available tRNA/aminoacyl-tRNA synthetase pairs orthogonal in mammalian cells, the pyrrolysyl tRNA/aminoacyl-tRNA synthetase pair from *Methanosarcina mazei* (*Mma* PylRS/PylT) being the most active and versatile to date. We found a previously uncharacterized pyrrolysyl tRNA/aminoacyl-tRNA synthetase pair from the human gut archaeon *Methanomethylophilus alvus Mx1201* (*Mx1201* PylRS/PylT) to be active and orthogonal in mammalian cells. We show that the new PylRS enzyme can be engineered to expand its ncAA substrate spectrum. We find that due to the large evolutionary distance of the two pairs, *Mx1201* PylRS/PylT is partially orthogonal to *Mma* PylRS/PylT. Through rational mutation of *Mx1201* PylT, we abolish its non-cognate interaction with *Mma* PylRS, creating two mutually orthogonal PylRS/PylT pairs. Combined in the same cell, we show that the two pairs can site-selectively introduce two different ncAAs in response to two distinct stop codons. Our work expands the repertoire of mutually orthogonal tools for genetic code expansion in mammalian cells and provides the basis for advanced *in vivo* protein engineering applications for cell biology and protein production.

## INTRODUCTION

Genetic code expansion allows for the addition of new chemical functionalities to proteins in living cells in the form of non-canonical amino acids (ncAAs). ncAAs are site-specifically installed through repurposing of a genetically encoded nonsense stop codon, most often *amber* (TAG). So-called *amber* suppression relies on introduction of a tRNA^CUA^/aminoacyl-tRNA synthetase pair into the cell that is orthogonal to, i.e. does not cross-react with, all endogenous tRNAs and aminoacyl-tRNA synthetases.

Nature created a repertoire of alternatives to the standard genetic code over billions of years of evolution. It is the rare outliers to the universal code that have provided useful molecular tools for synthetic biology (Baranov et al. 2015). The pyrrolysyl-tRNA (PylT)/pyrrolysyl-tRNA synthetase (PylRS) pair has become the most versatile tool for genetic code expansion in *E. coli*, yeast, mammalian cells and metazoan organisms. Pyrrolysine (Pyl, N-ε-4-methyl-pyrroline-5-carboxylate-L-lysine) is a biosynthetic amino acid, genetically encoded in a small number of methanogenic bacteria and archaea. In these organisms, a dedicated PylRS/PylT*^CUA^* pair directs Pyl incorporation in response to *amber* stop codons (Polycarpo et al. 2004). PylRS, the *PylS* gene product, accepts a range of structurally similar ncAAs in addition to its natural substrate. Further, PylRS has been a successful target for rational protein design and directed evolution, expanding the repertoire of accepted ncAA substrates (Wan et al. 2014). This includes ncAAs that carry chemical groups for bioorthogonal reactivity; photocaged amino acids for or photoactivated crosslinkers for photocontroled reactions; amino acids with native post-translational modifications (PTMs) (Chin 2017). The PylRS/PylT pair supports highly efficient recoding in mammalian cells (Elsässer et al. 2016;Schmied et al. 2014) enabling application of genetic code expansion technology to address biological questions in the context of the living cell.

In principle, reassignment of more than one natural codon could further augment the ability to engineer proteins harboring multiple ncAAs *in vivo*. Since PylT is not hardwired to recode amber codons, other stop codons can be recoded in mammalian cells (Ambrogelly et al. 2007;Xiao et al. 2013). Limiting to this application is the availability of mutually orthogonal tRNA/aminoacyl-tRNA synthetase pairs that are also orthogonal to the host cell. Thus, a key aim in the field is to find or rationally generate new mutually orthogonal pairs (Chatterjee et al. 2012;Neumann et al. 2010).

The two widely used PylRS/PylT pairs belong to the archaea *Methanosarcinales*, *M. mazei* (*Mma*) and *M. barkeri* (*Mba*), predominantly found in semi-aquatic environments. Recently, a number of new, evolutionary distant, Pyl-encoding archaea have been characterized from the human gut microbiome (Borrel et al. 2014;Borrel et al. 2012;Borrel et al. 2013;Dridi et al. 2012). Here, we explored the utility of the PylRS/PylT pair of *Candidatus Methanomethylophilus alvus Mx1201* (*Mx1201*) in mammalian cells. There were three rationales for this: First, we speculated that proteins of gut-resident archaea might have evolved to optimally perform at human body temperature as opposed to the environmental species that need to be adaptive over a wide temperature range (Gunnigle et al. 2013). Second, *Mx1201 PylS* gene encodes a smaller protein (31 kDa), which may be easier to express in a heterologous system. PylRS expressed in mammalian cells show predominantly nuclear localization which has been linked to inefficient function (Nikić et al. 2016). Third, divergent evolution from the *Methanosarcina* PylRS/PylT pairs could manifest in mutual orthogonality. Mutually orthogonal PylRS/PylT pairs would enable incorporation of two distinct non-canonical amino acids (ncAA) using a dedicated tRNA each, within the same host system.

## RESULTS AND DISCUSSION

### *Mx1201* PylRS/PylT pair has unique properties when compared to previously characterized archaeal and bacterial PylRS/PylT pairs

*Mx1201 PylS* encodes for a 275 amino acid protein, roughly half the size of *Mma* PylRS (Figure 1a). *Mx1201* PylRS is homologous to the C-terminal domain (CTD) of *Mma* PylRS only, and there is no gene product in the *Mx1201* genome with homology to the PylRS N-terminal domain (NTD) found in all previously characterized archaeal PylRS variants (Borrel et al. 2014). The PylRS CTD harbors the catalytic site, binding both Pyl and the anticodon stem of PylT. The NTD has been shown to bind the variable loop region on the opposite side of PylT (Figure 1b) and has been considered essential for aminoacylation activity *in vivo (Herring et al. 2007;Suzuki et al. 2017*). Notably, bacterial *PylS* genes encode two separate subunits PylSn and PylSc that structurally correspond to the two domains described above for archaea, suggesting that the complementing roles of PylRS CTD and NTD are conserved across the two kingdoms. In contrast to this, *Methanomethylophilus alvus Mx1201* and few related species were the first genomes to be discovered lacking any detectable PylSn homolog (Borrel et al. 2014), indicating that *Mx1201* PylRS may have evolved to function entirely without NTD. *Mx1201* PylRS and PylT show overall low sequence identity with the *Mma* PylRS/PylT pair (Figure 1a, b). *Mx1201* PylT is one of the most distant homologs of known archaeal PylT (Supplementary Figure 1), has a considerably divergent acceptor stem and appears to have an even further shortened D loop together with a ‘broken’ anticodon stem when compared to *Mma* PylT (Figure 1b). Given the many questions arising from such unusual features, we set out to characterize *Mx1201* activity, specificity and orthogonality in mammalian cells.

**Figure 1.**
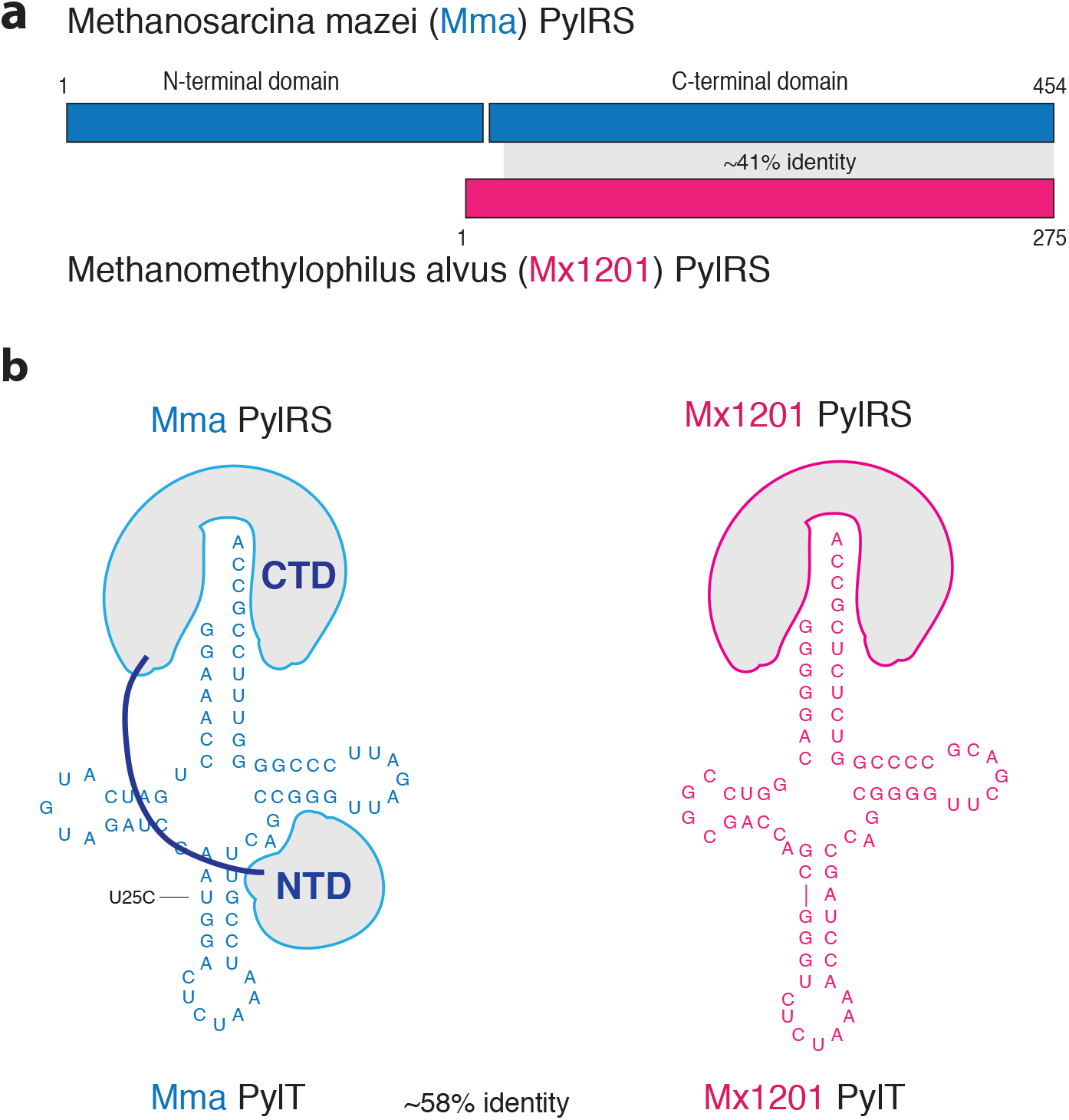
Divergent features of *Mx1201 and Mma* PylRS/PylT pairs. (a) *Methanosarcina mazei* (abbreviated as *Mma,* shown in blue) PylRS and *Methanomethylophilus alvus Mx1201* (abbreviated as *Mx1201*, shown in coral) PylRS domain structures and homology region. (b) Cloverleaf structure of *Mma* and *Mx1201* PylTs. Throughout this study, the previously described Mma PylT^U25C^ variant was used. Cartoons of cognate PylRS show sites of recognition between enzyme and tRNA.

### *Mx1201* PylRS/PylT is a functional amber suppressor, orthogonal to endogenous tRNAs and RS in mammalian cells

Previously, we have employed an efficient plasmid transfection system to direct ncAA incorporation into a GFP reporter protein in mammalian cells (Schmied et al. 2014;Elsässer 2018). Here, we cloned *Mma* and *Mx1201 PylS* and *PylT* genes into a similar plasmid design, expressing *PylS* from an EF1 promoter and 4x*PylT* from human U6 or 7SK promoters in tandem repeats (Supplementary Figure 2, Supplementary Table 1). For an initial combinatorial characterization, a plasmid expressing *Mma* or *Mx1201 PylS* was cotransfected with a second plasmid expressing four copies of either PylT variant together with the sfGFP^150TAG^ reporter. The *Mma PylT* used in this study has a base substitution in the anticodon stem, U25C, previously found to increase amber suppression efficiency in *E. coli* and mammalian cells (Chatterjee et al. 2013;Schmied et al. 2014). Transient transfection was performed in human embryonic kidney (HEK293T) cells. Amber suppression was measured by GFP fluorescence in cell lysates and by western blotting. We used cyclopropene-L-Lysine (CpK, Supplementary Figure 3) as ncAA, which is efficiently incorporated with wildtype *Mma* PylRS/PylT (Schmied et al. 2014;Elliott et al. 2014).

First, we sought to test if the *Mx1201* PylRS/PylT pair was functional in mammalian cells. Expression of *Mx1201* PylRS/PylT pair indeed allowed selective incorporation of CpK into the GFP reporter, with 28% yield as compared to a no-stop GFP control (Figure 2a, b; Supplementary Figure 4a). In comparison, the *Mma* PylRS/PylT pair reached 93% (Figure 2a).

**Figure 2.**
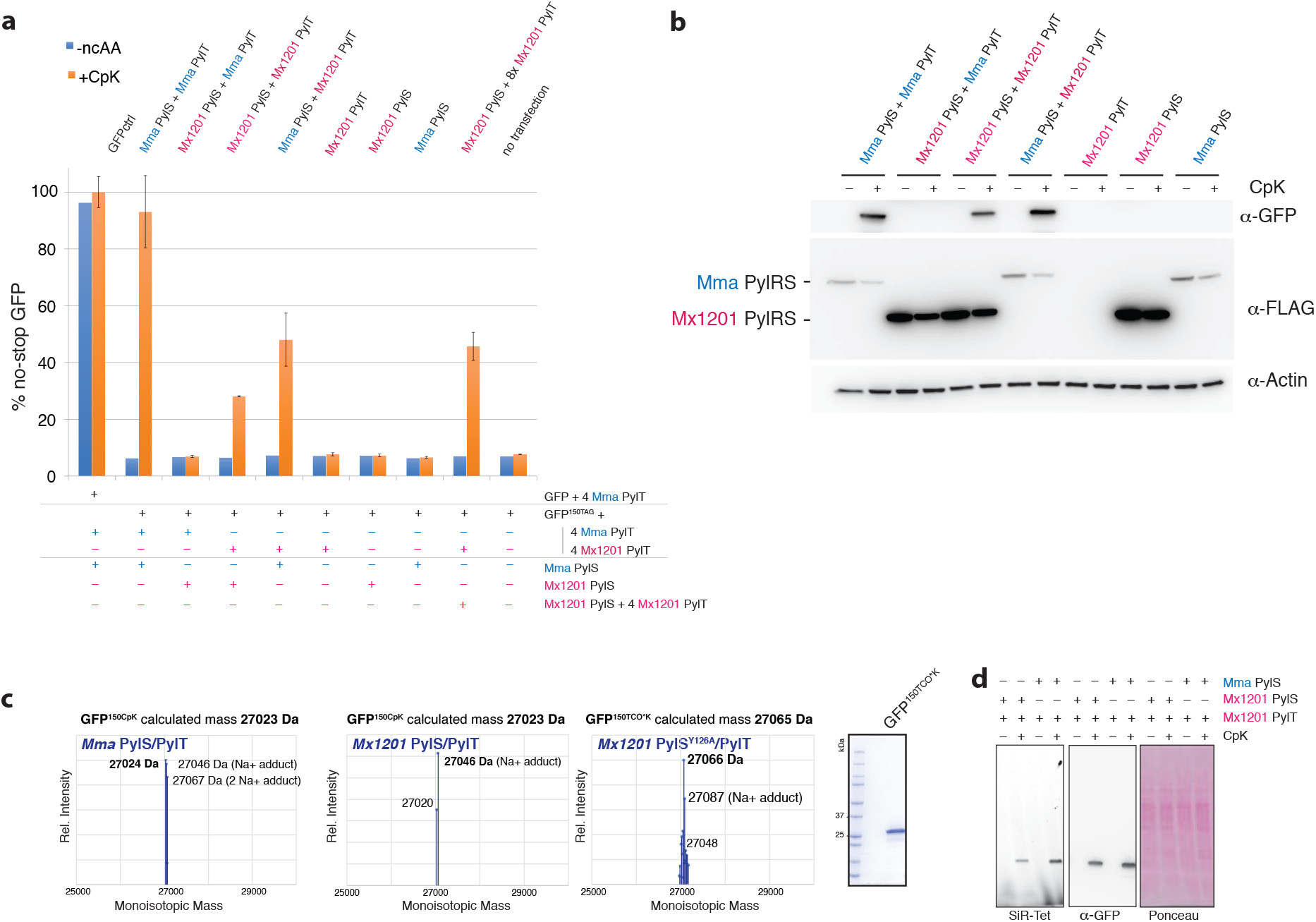
*Mx1201* PylRS/PylT pair is active and orthogonal in mammalian cells; *Mx1201* PylRS *and Mma* PylT are orthogonal whereas *Mma* PylRS charges both *Mma* and *Mx1201* PylT. (a) Fluorescence plate reader assay from HEK293T cell lysates transiently transfected with a GFP^150TAG^ reporter, and a combination of tRNA and synthetase, at a 9:1 ratio. GFP fluorescence is shown as percentage of fluorescence measured with a GFP construct without TAG stop codon (GFPctrl) in the same experiment. For each combination, quadruplicate transfections were performed. For three of the four samples, medium was supplemented with 0.2 mM CpK, all samples were harvested 24h post transfection. See Supplementary Figure 4 for fluorescence microscopy pictures of transfected cells. (b) Western blot from HEK293T cell lysates transiently transfected with the reporter GFP^150TAG^ reporter, and a combination of tRNA and synthetase, showing the expression of GFP, FLAG-tagged synthetase variants, and a β-actin loading control. (c) Intact mass determination of purified GFP containing CpK at position 150 produced with *Mma* and *Mx1201* PylRS/PylT, as well as TCO^*^K-containing GFP produced with *Mx1201* PylRS^Y126A^/PylT. All deconvoluted monoisotopic masses in the 25-30kDa mass range are graphed, and the predicted monoisotopic mass is given for comparison. (d) In-gel far-red fluorescence image and western blot against GFP from HEK293T cell lysates. Lysates have been labeled with silicon rhodamine (SiR) tetrazine (SiR-Tet) fluorescent dye.

The PylRS/PylT pairs encoded by *Mx1201* and *Mma* differ substantially in their primary sequences. We sought to understand if these PylRS and PylT would cross-react or were in fact non-functional accross species, thus mutually orthogonal. Interestingly, *Mma* PylRS aminoacylates *Mx1201* PylT more efficiently (48% suppression) than *Mx1201* PylRS, suggesting optimal enzymatic activity even for the non-cognate PylT. This result also implies that key structural recognition features are conserved between the two distant PylT relatives. In contrary, *Mx1201* PylRS did not elicit any measurable amber suppression with *Mma* PylT. Western blotting and immunostaining using an N-terminal FLAG-tag confirmed expression of *Mx1201* PylRS (Figure 2a). *Mx1201* PylRS protein levels appeared much higher than for *Mma* PylRS in our lysates (Figure 2a).

However, it should be noted that *Mma* PylRS has a distinctive nuclear localization (Supplementary Figure 6), and we only solubilized 50% of the *Mma* PylRS protein using RIPA buffer (Supplementary Figure 4b). *Mx1201* PylRS is soluble and mostly cytosolic (Supplementary Figure 4b, 6). In summary, *Mx1201* PylRS is stable and correctly localized in mammalian cells, but does not generate aminoacylation activity equivalent to *Mma* PylRS. Given the known importance of the PylRS NTD for PylT binding (Herring et al. 2007;Suzuki et al. 2017), we speculate that the *Mx1201* PylRS evolved means to partially but not completely compensate for the lack of an NTD.

Further, we tested if additional *PylT* copies would enhance activity of the *Mx1201* PylRS/PylT pair. Indeed, supplying 4x*Mx1201 PylT* on both plasmids raised the amber suppression efficiency to 46% of a no-stop GFP (Figure 2a).

To confirm orthogonality of *Mx1201* PylRS/PylT in mammalian cells, we further needed to ensure that *Mx1201* PylT is not charged by an endogenous aminoacyl-tRNA synthetase, and that *Mx1201* PylRS does not charge other tRNAs with CpK. sfGFP^150TAG^ expression was undetectable in the absence of CpK as judged by fluorescence measurement or western blot (Figure 2a, b). Further, we determined the identity of CpK incorporated into the sfGFP^150TAG^ construct by mass spectrometry (Figure 2c). *Vice versa*, *Mx1201* PylRS does not charge other mammalian tRNAs, as shown by selective SiR-tetrazine reaction with the amber-suppressed GFP and the absence of additional labeled endogenous proteins in whole-cell lysates (Figure 2d).

Together, these results show that the *Mx1201* PylRS/PylT pair is functional and orthogonal in mammalian cells.

### *Mx1201* PylRS can be engineered for expanded substrate specificity

The key advantage of the PylRS/PylT system in mammalian cells over other orthogonal tRNA systems lies in the ability to incorporate structurally diverse ncAAs with useful functions for probing, controlling and engineering proteins in living cells. This is both facilitated by the relative promiscuity of the wildtype PylRS enzyme and the ability to generate active site mutations with expanded substrate repertoire. Exploring the substrate preferences of PylRS/PylT pairs from additional species may further expand the available repertoire of PylRS variants. Having established a new PylRS/PylT pair in mammalian cells, we sought to understand the substrate specificity of *Mx1201* PylRS on a small set of functionally interesting ncAAs for mammalian cell biology (Supplementary Figure 3). While *Mma* PylRS accepted CpK as well as 3’-azibutyl-N-carbamoyl-lysine (AbK) (Figure 3a), *Mx1201* PylRS did not incorporate any of the tested ncAAs except CpK (Figure 3b).

**Figure 3.**
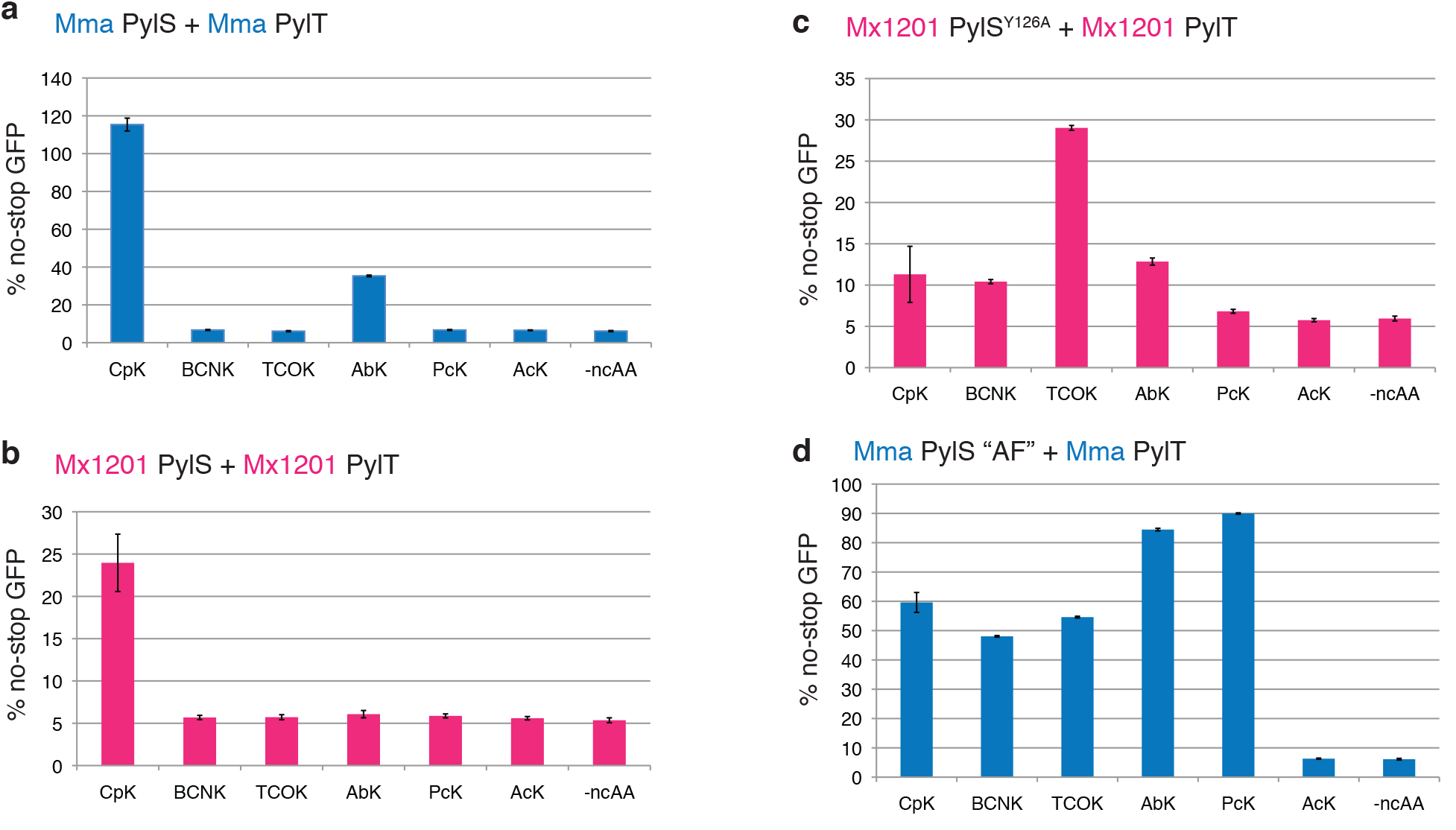
Substrate specificities of *Mx1201* PylRS/PylT and *Mx1201* PylRS^Y126A^/PylT. Fluorescence plate reader assay from HEK293T cell lysates transiently transfected in 4:1 ratio with a GFP^150TAG^ reporter and (a) *Mma* PylRS/PylT, (b) *Mx1201* PylRS/PylT, (c) *Mx1201* PylRS^Y126A^/PylT or (d) *Mma* PylRS AF/PylT. GFP fluorescence is shown as percentage of fluorescence measured with a GFP construct without TAG stop codon in the same experiment. Cells were grown for 24 hours in the absence (-ncAA) or presence of one of the following ncAAs: 0.2 mM CpK, 0.5 mM BCNK, 0.1 mM TCO^*^K, 0.5 mM AbK, 0.5 mM PcK, 10 mM AcK. For full names and structures of amino acids refer to Supplementary Figure 3.

Sequence alignment and modeling suggests that, despite overall low sequence identity, the Pyl-binding pocket of *Mx1201* PylRS is highly similar to other archaeal and bacterial PylRS homologs (Supplementary Figure 5a). Few exceptions apply, such as Met129 and Val168 at the distal end contacting the pyrrole ring, where most other PylRS, including *Mma* PylRS, feature a highly conserved Leu and Cys residue, respectively (Supplementary Figure 5b). Thus, there may be subtle variations in the substrate binding pocket underlying the more restricted repertoire of *Mx1201* PylRS.

To expand the scope of the *Mx1201* PylRS/PylT pair in mammalian cells, we tested the possibility of engineering the *Mx1201* PylRS substrate binding pocket. We generated a variant, *Mx1201* PylRS^Y126A^, corresponding to a *Mma* PylRS Y306A mutant. The latter residue caps the Pyl-binding pocket in available PylRS structures and has previously been described to limit PylRS from incorporating ncAAs with longer and/or larger side chains than Pyl (Yanagisawa et al. 2008;Kavran et al. 2007). *Mx1201* PylRS^Y126A^ has a much reduced activity towards CpK, but intriguingly gained activity (yield 29% of no-stop GFP) for axial trans-Cyclooct-2-ene–Lysine (TCO^*^K). Comparing this result with the prior PylRS variant described for TCO^*^K, PylRS-AF (Nikić et al. 2014;Yanagisawa et al. 2008), *Mx1201* PylRS^Y126A^ has roughly half the activity but dramatically increased specificity for TCO^*^K over other ncAAs tested.

### *Mma* PylRS requires its NTD for activity towards *Mx1201* PylT

Following up on our finding that *Mma* PylRS accepts *Mx1201* PylT (Figure 1a, b), we sought to understand how *Mma* PylRS recognizes the non-cognate PylT. Specifically, we wondered if *Mma* PylRS N- and C-terminal domains differentially contributed to cognate *Mma* PylT or non-cognate *Mx1201* PylT binding. We created a new construct, *Mma* PylRS-CTD, by deleting the first 187 residues of *Mma* PylRS (Figure 4a). This fragment contains the entire region homologous to the *Mx1201* PylRS enzyme (see Figure 1a).

**Figure 4.**
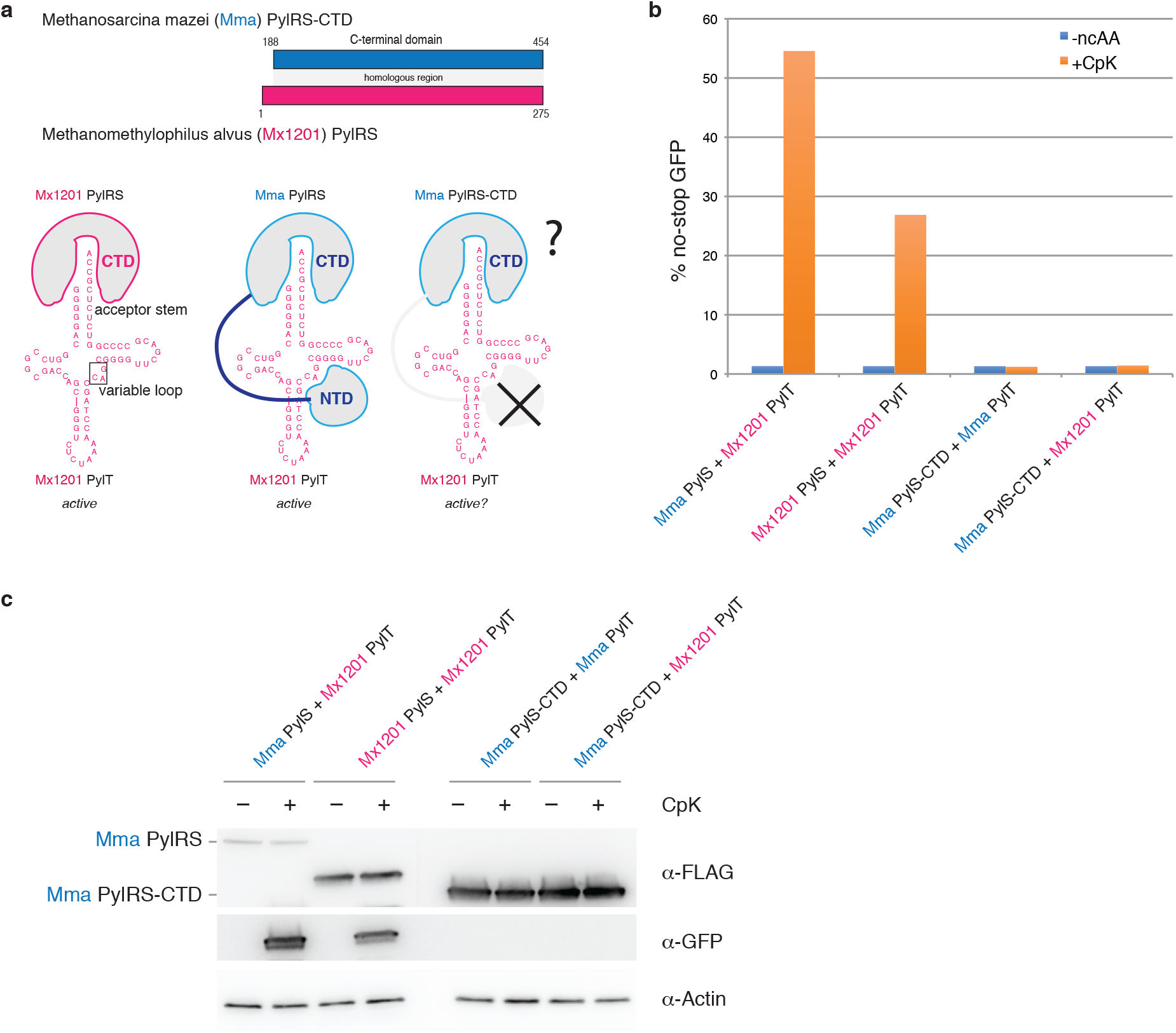
*Mma* PylRS interacts with non-cognate *Mx1201* PylT through its N-terminal domain. (a) Top scheme shows the C-terminal domain (CTD) construct used for *Mma* PylRS, containing the region corresponding to full-length *Mx1201* PylRS. Bottom scheme indicates known identity elements (acceptor stem and variable loop) on putative *Mx1201* PylT structure and putative binding regions for *Mx1201* PylRS and *Mma* PylRS. (b) Fluorescence plate reader assay from HEK293T cell lysates transiently transfected in 9:1 ratio with a GFP^150TAG^ reporter and the indicated combination of tRNA and synthetase. GFP fluorescence is shown as percentage of fluorescence measured with a GFP construct without TAG stop codon in the same experiment. Cells were grown in the presence or absence of 0.2 mM CpK for 48h. (c) Western blot showing the expression of FLAG-tagged synthetase variants, GFP and a β-actin loading control.

*Mma* PylRS-CTD was inactive with its cognate PylT (Figure 4b, c). Essentiality of the *Mma* PylRS NTD has never been explicitly tested in mammalian cells but mirrors observations made in *E. coli* and *in vitro* (Herring et al. 2007). Unlike the full-length enzyme, *Mma* PylRS-CTD also did not elicit measurable activity with *Mx1201* PylT (Figure 4b, c). These results show that *Mma* PylRS activity towards *Mx1201* PylT is fully dependent on its N-terminal domain. The results imply two fundamentally different binding modes employed by *Mma* and *Mx1201* PylRS: *Mx1201* PylRS has evolved efficient tRNA binding through its catalytic domain, making an N-terminal domain obsolete. This binding mode must be facilitated by specific features of the cognate *Mx1201* PylT, since *Mx1201* does not accept *Mma* PylT (Figure 1a). *Mma* PylRS, on the other hand, employs the canonical NTD interaction with the variable loop region of PylT, a known identity element of PylRS/PylT pairs (Jiang and Krzycki 2012;Ambrogelly et al. 2007). Supporting this hypothesis, the variable loop itself (^41^CAG^43^) and adjacent bases of *Mx1201* PylT are conserved to their *Mma* PylT counterparts (Figure 4a).

### Generation of an orthogonal PylRS/PylT pair by disrupting NTD interaction of *Mx1201* PylT

Above findings provide key prerequisites for the creation of mutually orthogonal PylRS/PylT pairs in mammalian cells: Two evolutionary distant PylRS enzymes (and engineered variants) with partially, but not fully overlapping, substrate specificities, that use distinct surfaces for their recognition of the respective cognate PylT. We hypothesized that rationally designed mutations in the variable loop region of *Mx1201* PylT would directly interfere with the non-cognate recognition by *Mma* PylRS NTD (Figure 5a). Disrupting this interaction would make the *Mx1201* and *Mma* tRNA/aminoacyl-tRNA synthetase pairs mutually orthogonal.

**Figure 5.**
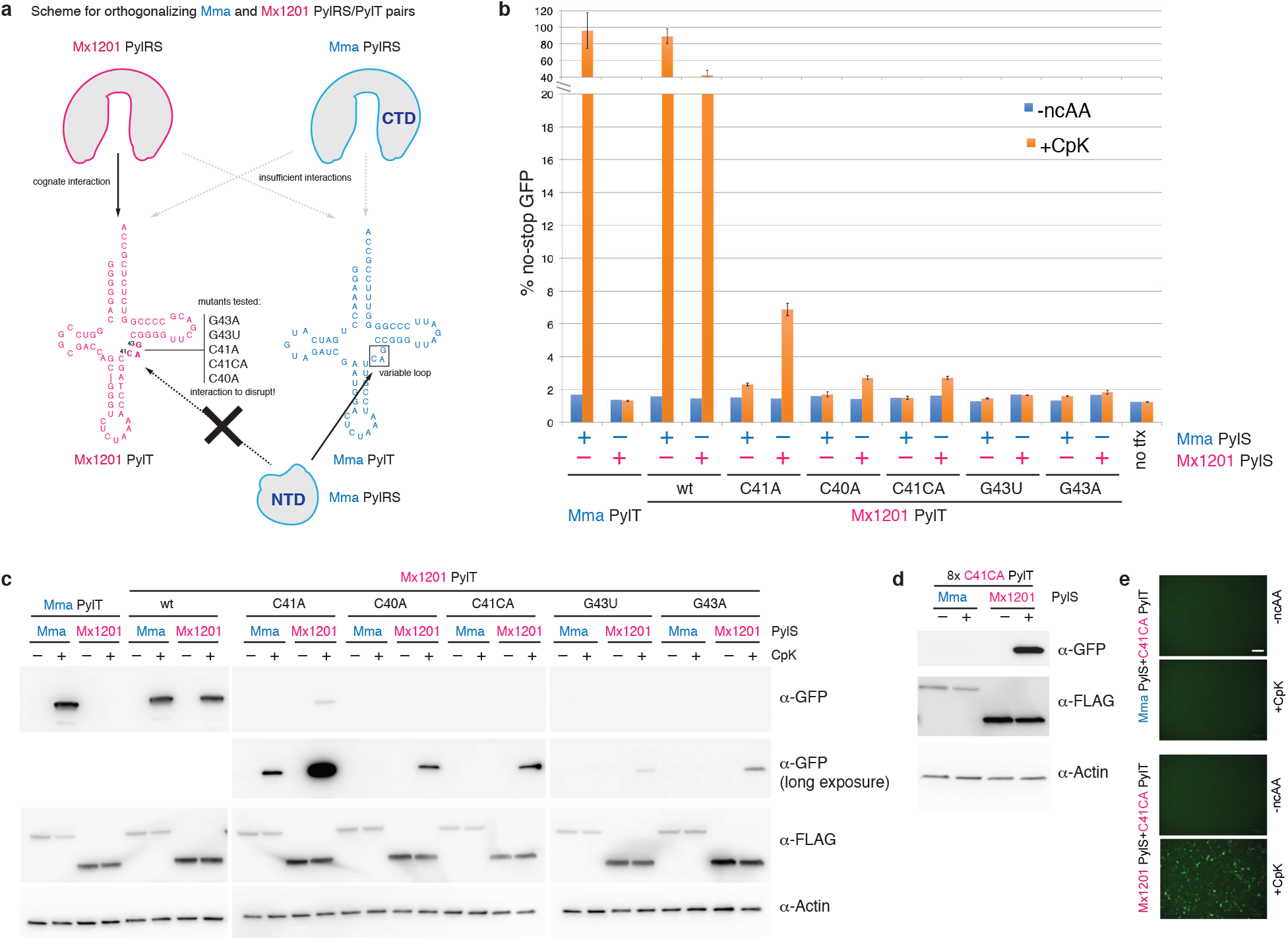
Mutations in the *Mx1201* PylT variable loop disrupt recognition by *Mma* PylRS creating orthogonal PylRS/PylT pairs. (a) Scheme depicting predominant modes of recognition between PylRS and cognate PylTs: *Mx1201* PylRS, in the absence of an NTD, must bind PylT through the acceptor stem only. Our results show that *Mma* PylRS alone recognizes neither *Mma* PylT nor *Mx1201* PylT in the absence of its NTD, indicating that the recognition of the acceptor stem via its CTD is secondary to engagement of its NTD with the variable loop. Further indicated are mutations introduced in the variable loop of *Mx1201* PylT that are predicted to abolish interaction with the *Mma* PylRS NTD. (b) Fluorescence plate reader assay of HEK293T cell lysates transiently transfected in 5:1:4 ratio with a GFP^150TAG^ reporter and the indicated synthetase and tRNA. GFP fluorescence is shown as percentage of fluorescence measured with a GFP construct without TAG stop codon in the same experiment. For each combination, quadruplicate transfections were performed. For three of the four samples medium was supplemented with 0.2 mM CpK, all samples were harvested 48h post transfection. Note the broken Y axis. (c) Western blot showing the expression of FLAG-tagged synthetase variants, GFP and a β-actin loading control. (d) Western blot of HEK293T cell lysates transiently transfected in 1:9 ratio of GFP^150TAG^ reporter and either *Mx1201* PylT^C41CA^/*Mma* PylRS or *Mx1201* PylT^C41CA^/*Mx1201* PylRS, showing the expression of FLAG-tagged synthetase variants, GFP and a β-actin loading control. CpK was added at the time of transfection, samples were harvested 48h post transfection. (e) Fluorescent images of transfected HEK293T cells used for panel (d) prior to cell lysis. White scale bar is 100μm.

A recent crystal structure of the PylRS-NTD–PylT complex reveals that the recognition relies predominantly on steric constraints that would exclude any more spacious variable loop (Suzuki et al. 2017). We reasoned that introduction of variations to the variable loop analogous to those shown to abrogate PylRS binding and aminoacylation (Jiang and Krzycki 2012;Ambrogelly et al. 2007), would provide candidates for disrupting the interaction between *Mx1201* PylT and *Mma* PylRS.

As a caveat to this simple approach, even isolated base changes may affect folding of the tRNA in unexpected ways, particularly since the short variable loop participates in a tightly packed tertiary core (Nozawa et al. 2009). We chose to test four single base substitutions, C40A, C41A, G43A, G43U, and one insertion C41CA (Figure 5a).

For a more facile generation and screening of *Mx1201* PylT mutants, we moved to a three-plasmid expression system where PylRS and sfGFP^150TAG^ reporter were expressed from plasmids without *PylT*, and *PylT* was supplied on a third plasmid (Supplementary Figure 2). Wild type and mutant *Mx1201* PylT were coexpressed with the same sfGFP^150TAG^ reporter and either *Mma* PylRS or *Mx1201* PylRS (Figure 5b, c). As in previous experiments, *Mx1201* PylT was more active with *Mma* PylRS (89% of no-stop GFP) than its cognate *Mx1201* PylRS (42% of no-stop GFP).

None of the *Mx1201* PylT mutants exhibited a similar efficiency, suggesting that mutations in the variable loop affected both cognate and non-cognate interactions with *Mx1201* PylRS and *Mma* PylRS, respectively. Nevertheless, C40A, C41A and C41CA mutations preserved measurable activity with *Mx1201* PylRS. Of these, C41A retained the highest activity with *Mx1201* PylRS (7% of no-stop GFP) while disproportionally reducing *Mma* PylRS activity (Figure 5b, c). Since we aimed to create a fully orthogonal pair, we focused on C41CA, which showed lower activity with *Mx1201* PylRS but lost all detectable activity with *Mma* PylRS even by sensitive western blot (Figure 5c). Moving the *Mx1201* PylT^C41CA^ mutant back to our efficient two-plasmid transfection system (Supplementary Figure 2), we were able create robust amber suppression with *Mx1201* PylRS, and confirm orthogonality to *Mma* PylRS (Figure 5d, e; for full panel see Supplementary Figure 7).

In conclusion, we we have created a new tRNA/aminoacyl-tRNA synthetase pair, *Mx1201* PylS/*Mx1201* PylT^C41CA^, that is orthogonal in mammalian cells, and also mutually orthogonal to the *Mma* PylRS/PylT pair.

### Site-specific incorporation of two distinct ncAAs using two orthogonal PylRS/PylT pairs

Next, we aimed to implement dual site-specific protein modification at independent sites and with distinct ncAAs in mammalian cells using the two orthogonal PylRS/PylT pairs. This first required to further modify one pair to recode another stop codon. We chose *ochre* (TAA) because it is only marginally more abundant than *amber* in mammalian cells. We thus created *Mma* PylT^UUA^, where the anticodon was changed from CUA to UUA. We used a fluorescent reporter, sfGFP^102TAG+150TAA^, containing two stop codons, 102TAG and 150TAA.

To incorporate two distinct ncAAs, the two orthogonal PylRS must each incorporate one of the ncAAs with high selectivity (Figure 6a). Since we have shown that wild type *Mx1201* PylRS and the engineered variant PylRS^Y126A^ both have narrow substrate specificities, they can be combined in many unique combinations with known *Mma* PylRS variants evolved for specific ncAAs, e.g. photocaged lysine (PcK), acetyl-lysine (AcK), diazirine lysine (AbK) (Neumann et al. 2008;Gautier et al. 2010;Ai et al. 2011).

**Figure 6.**
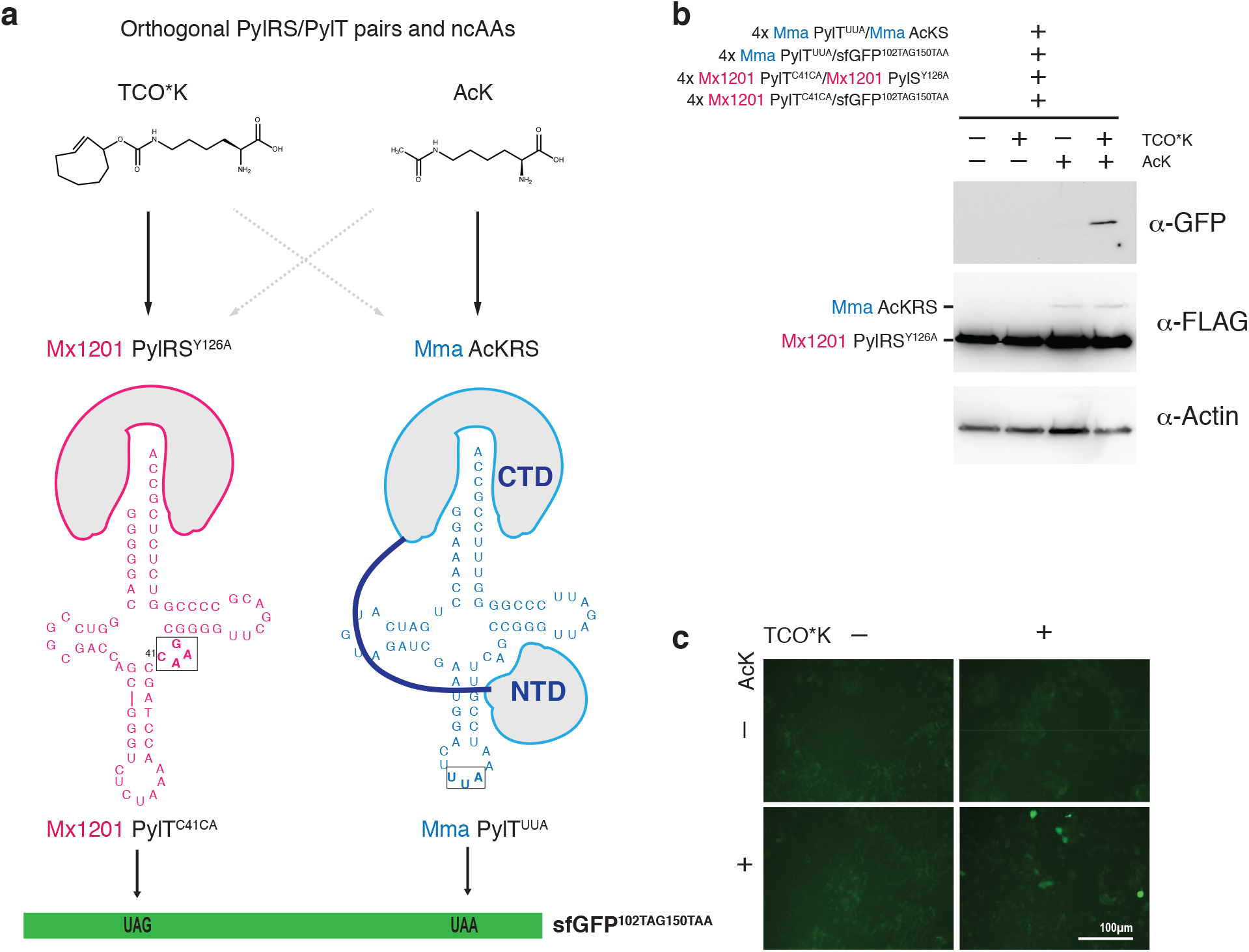
Dual ncAA incorporation in mammalian cells using mutually orthogonal PylRS/PylT pairs. (a) Scheme for dual ncAA incorporation using mutually orthogonal PylRS/PylT pairs in mammalian cells. Two PylRS variants with narrow substrate specificity were chosen to incorporate two different ncAAs (TCO^*^K and AcK) in response to distinct stop codons. Chemical structures of ncAA are given. PylRS and PylT variants used in this experiment are indicated. (b) Western blot of HEK293T cells transiently transfected in 4:4:1:1 ratio with the following components: GFP^102TAG150TAA^ reporter with 4x *Mx1201* PylT^C41CA^; GFP^102TAG150TAA^ reporter with 4x *Mma* PylT^TAA^; *Mx1201* PylRS^Y*126A*^ with 4x *Mx1201* PylT^C41CA^; *Mma* AcKRS with 4x *Mma* PylT^TAA^. Cells were grown in the presence of either none, 0.1 mM TCO*K, 10 mM AcK or both ncAAs for 48h. Lysates were analyzed for expression of FLAG-tagged synthetase variants, GFP and a β-actin loading control. Notably, detectable AcKRS protein levels are dependent on the presence of AcK substrate (Figure 6b). (c) Fluorescence images of transfected HEK293T cells used for panel (b) prior to cell lysis.

Here, we chose to incorporate AcK and TCO^*^K into the same protein, GFP, in response to TAA and TAG stop codons, respectively: we created a pair with *Mma* PylT^UUA^ and AcKRS (Neumann et al. 2008) and combined it with the *Mx1201* PylRS^Y126A^/*Mx1201* PylT^C41CA^ pair in the same cell (Figure 6a). Indeed, we were able to observe production of GFP in cells transfected with the two PylRS/PylT pairs and the sfGFP^102TAG+150TAA^ reporter in the presence of both ncAA, but not with either individually or none (Figure 6b, c). Thus, we have shown that the two orthogonal PylRS/PylT pairs can be employed together in the same cell to site-specifically incorporate two distinct ncAAs.

## CONCLUSIONS

We have set out to identify new orthogonal tRNA/aminoacyl-tRNA synthetase pairs in mammalian cells. Exploring the function of a previously uncharacterized distant homolog of the *Mma* PylRS/PylT pair from *Methanomethylophilus alvus Mx1201*, we found that the *Mx1201* PylRS/PylT pair is active and orthogonal in mammalian cells. *Mx1201* PylRS is also orthogonal to *Mma* PylT. We show that the *Mx1201* PylRS enzyme can be engineered to expand its ncAA substrate spectrum, creating a new variant *Mx1201* PylRS^Y126A^ that can efficiently incorporate TCO^*^K. *Mx1201* PylRS is a small enzyme (31 kDa) that lacks the N-terminal domain present in all previously characterized archaeal PylRS proteins. *Mma* PylRS can accept the non-cognate *Mx1201* PylT as substrate, and intriguingly we find that this is entirely dependent on its NTD. This indicated that *Mx1201* and *Mma* PylRS rely on different identity determinants of PylT. We used this knowledge to rationally design a series of mutant *Mx1201* PylT with modified variable loop region in order to selectively disrupt its interaction with *Mma* PylRS. We identify several *Mx1201* PylT mutants that retain their cognate interaction with *Mx1201* PylRS but reduce or abrogate non-cognate recognition by *Mma* PylRS. We selected one mutant, *Mx1201* PylT^C41CA^ for further experiments. We show that the *Mx1201* PylS/ PylT^C41CA^ pair is orthogonal to *Mma* PylRS/PylT. Combined in the same cell, we show that the two pairs can introduce two different ncAAs in response to two distinct stop codons. Our findings shed light into a new clade of PylRS enzymes with unexpected properties, functionally divergent from the previously studied archaeal and bacterial system. Our study agrees with a recent first characterization of the *Mx1201* PylRS/PylT pair in *E. coli* published during the preparation of this manuscript (Willis and Chin 2018). Notably, *Mx1201* PylRS/PylT pair exceeds the activity of *Mma* PylRS/PylT pair in *E.coli*. Chin and Willis also used a similar rational for creating orthogonal PylRS/PylT pairs and showed their excellent activity in *E.coli*. Intriguingly their directed evolution approach to identify most highly active orthogonal *Mx1201* PylTs yields additional candidates for mammalian cells. Our work expands the repertoire of mutually orthogonal tools for genetic code expansion in mammalian cells and provides the basis for advanced *in vivo* protein engineering applications for cell biology and protein production.

## MATERIALS AND METHODS

### DNA constructs

The sfGFP 150L control reporter construct has been described previously (Schmied 2014). A series of plasmids for amber suppression (pAS) was created based on PB510B-1 (System Biosciences) piggybac plasmid (Table 1 and Supplementary Figure 2). pUC-based plasmids were used to express tRNA from single copy genes (Table 2 and Supplementary Figure 2). All DNA constructs were verified by Sanger sequencing.

**Table 1.** pAS plasmids for EF1α controlled expression of *PylS* and reporter genes.

| plasmid number | 4xtRNA cassette | EF1α promoter |
| --- | --- | --- |
| 1 | – | FLAG_Mma PyIRS |
| 2 | 7SK-Mma PyIT | FLAG_Mma PyIRS |
| 3 | 7SK-Mma PyIT | FLAG_Mma PyIRS/AF |
| 4 | 7SK-Mma PyIT | FLAG_Mma PyIRS CTD |
| 5 | 7SK-Mx1201 PyIT C41CA | FLAG_Mma PyIRS |
| 6 | 7SK-Mma PyIT<sup>UUA</sup> | FLAG_AcKRS |
| 7 | – | FLAG_Mx1201 PyIRS |
| 8 | U6-Mx1201 PyIT | FLAG_Mx1201 PyIRS |
| 9 | U6-Mx1201 PyIT | FLAG_Mx1201 PyIRS Y126A |
| 10 | 7SK-Mx1201 PyIT | FLAG_Mx1201 PyIRS |
| 11 | 7SK-Mx1201 PyIT | FLAG_Mx1201 PyIRS Y126A |
| 12 | 7SK-Mx1201 PyIT C41CA | FLAG_Mx1201 PyIRS |
| 13 | 7SK-Mx1201 PyIT C41CA | FLAG_Mx1201 PyIRS Y126A |
| 14 | – | sfGFP 150TAG |
| 15 | 7SK-Mma PyIT | sfGFP 150TAG |
| 16 | 7SK-Mma PyIT<sup>UUA</sup> | sfGFP 102TAG 150TAA |
| 17 | U6-Mx1201 PyIT | sfGFP 150TAG |
| 18 | 7SK-Mx1201 PyIT | sfGFP 150TAG |
| 19 | 7SK-Mx1201 PyIT C41CA | sfGFP 150TAG |
| 20 | 7SK-Mx1201 PyIT C41CA | sfGFP 102TAG 150TAA |

**Table 2.** PylT expression plasmids are pUC based.

| plasmid number | single copy tRNA gene |
| --- | --- |
| 21 | 7SK-Mma PyIT |
| 22 | 7SK-Mx1201 PyIT |
| 23 | 7SK-Mx1201 PyIT C41CA |
| 24 | 7SK-Mx1201 PyIT G43U |
| 25 | 7SK-Mx1201 PyIT C40A |
| 26 | 7SK-Mx1201 PyIT G43A |
| 27 | 7SK-Mx1201 PyIT C41A |

**Table 3.**
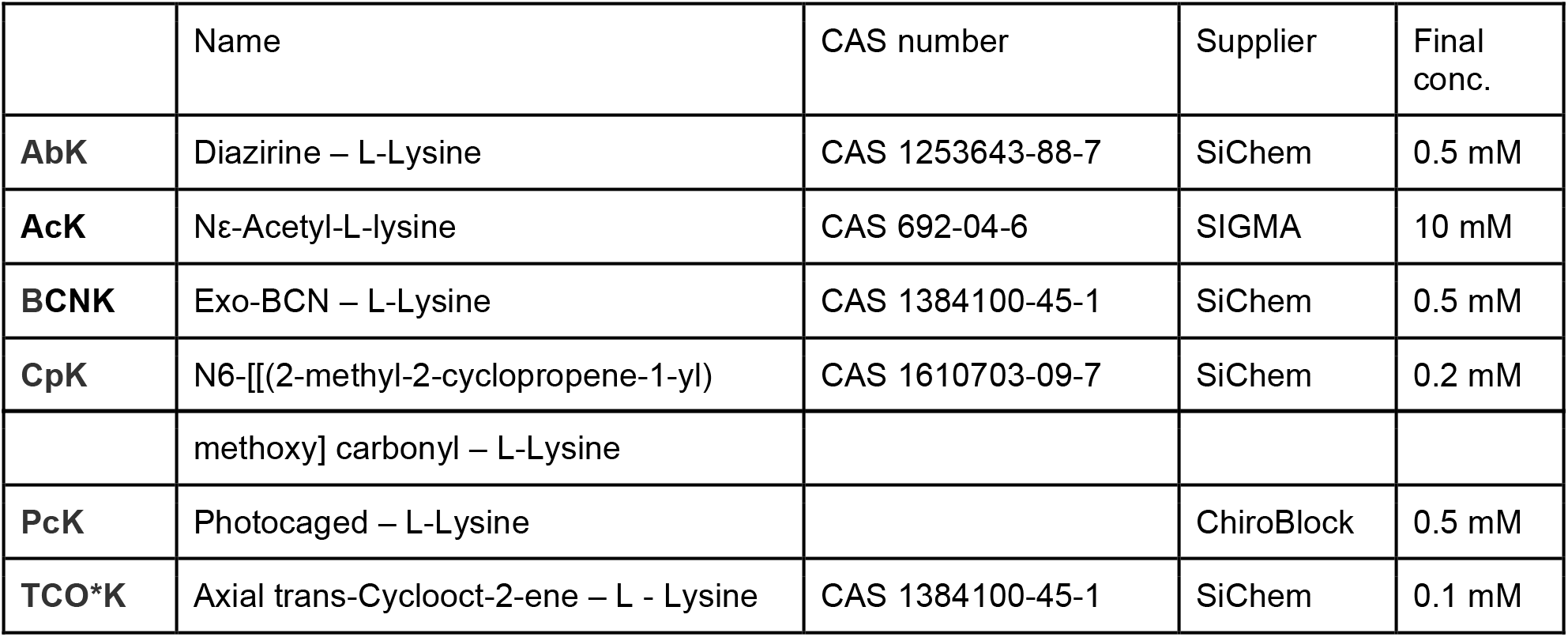
non-coding amino acids used in this study.

### Cell culture and Transfection

HEK293T cells were maintained in adherent culture at 37°C and 5 %CO_2_ atmosphere in Dulbecco’s modified Eagle’s medium (DMEM) (AqDMEM^TM^ high glucose, Sigma) supplemented with 10% FBS. For transfection 1.5 - 2.0×10^5^ HEK293T cells were seeded per well in 24-well plates 1d before transfection. Cells were transiently transfected with TransIT®-LT1 (Mirus) following manufacturer’s instructions. ncAAs were added at the time of transfection as indicated and transfected cells harvested 24h or 48h post transfection.

### Amino acids

Non-canonical amino acids (ncAAs) used in this study are summarized below. To incorporate ncAA into proteins, amino acid working solutions were prepared from 100 mM stock solutions and added to the cultured cells together with the transfection mixture.

### Mass Spectrometry

HEK293T cells were transfected and cultured in presence of ncAA for 72h. After cell lysis in RIPA buffer with added cOmplete protease inhibitor (Roche), the insoluble fraction was removed by centrifugation. GFP was captured on GFP-Trap_M magnetic beads (ChromoTek) and eluted with 1% acetic acid.

Mass spectrometric analysis was carried out at the Proteomics Biomedicum core facility, Karolinska Institutet, Stockholm. Intact protein mass analysis was performed using a TriVersa NanoMate chip-based electrospray device (Advion, Ithaca, NY) coupled to the LTQ Velos Orbitrap Elite (Thermo Scientific, Bremen, Germany). The ChipSoft Manager software was used to control the TriVersa NanoMate, while data was acquired directly from the Tune software of the mass spectrometer. The NanoMate delivered 2 μL of sample solution to the tip engaged with the back of the ESI chip and nano-spray ionization was initiated applying 1.9 kV and 0.8 psi gas pressure. The mass spectrometer was operated in positive ion mode with activated protein mode settings. MS data was collected in full scan mode (m/z 500-2000) with a resolution of 100,000 at m/z 400. Each scan comprises 1 microscan. The mass spectra shown are comprised of approximately 20 scans. Automatic gain control (AGC) was used to accumulate sufficient ions for analysis targeting 3×10^7^ ions in a maximum fill time of 200 ms. Data were analyzed using the Protein Deconvolution v3.0 software (Thermo Scientific) to calculate monoisotopic masses.

### Quantification of GFP expression

Transfected HEK293T cells were grown in presence of ncAA as indicated for 24h or 48h. Cells were lysed in RIPA buffer with 1× cOmplete protease inhibitor (Roche), the insoluble fraction was removed by centrifugation. GFP bottom fluorescence of an aliquot was measured in Tecan Infinity M200 pro platereader (excitation 485 nm, emission 518 nm). Fluorescence measurements were normalized to total protein content of each sample as determined by Pierce BCA assay kit (Fisher Scientific) on the same sample.

### Live-cell imaging for GFP expression

GFP expression was visualized by imaging in a ZOE™ Fluorescent Cell Imager (BioRad).

### Western Blot

Equal amounts of protein, as determined by Pierce BCA assay (Fisher Scientific) were separated on 4-20% Tris-glycine gels (BioRad) and transferred to nitrocellulose membranes. Expression of the sfGFP reporter and aminoacyl-tRNA synthetases was confirmed by immunoblotting with antibodies against GFP (Santa Cruz, sc-9996), FLAG-HRP (Sigma, A8592), β-Actin (cell signaling) and corresponding secondary HRP-conjugated antibodies when needed (BioRad).

### Immunostaining and fluorescence microscopy

Transfected cells were cultured in presence of ncAA for 24h. Before fixation (4% formaldehyde) and permeabilization (0.1% triton) the ncAA was removed for 8h. Samples were blocked in 2% BSA in TBS-T and subsequently incubated in presence of 0.5 mM SiR-tetrazine (Spirochrome). After washing with TBS-T samples were incubated with primary antibodies mouse anti-GFP (B-2, Santa Cruz #9996) and rabbit anti-FLAG (D6W5B, Cell Signaling #14793) and subsequently incubated with secondary antibodies anti-mouse Alexa Fluor 488, anti-rabbit Alexa Fluor 555 (Life Technologies) and DAPI (Sigma-Aldrich). After washing, cells were imaged on a Nikon eclipse Ti2 inverted widefield microscope, using a 20× 0.75 NA objective.

## Supplementary data availability

Plasmid sequences are deposited on Mendeley Data (<u>doi:10.17632/bnm4×5vjrs.1</u>).

## ACKNOWLEDGEMENTS

Mass Spectrometry service was provided by Proteomics Biomedicum core facility, Karolinska Institutet, Stockholm. We thank Akos Vegvari for data analysis and advice.

We thank the Jiri Bartek lab for access to Tecan Infinity M200 Pro plate reader and Nikon eclipse Ti2 microscope.

## AUTHOR CONTRIBUTIONS

SJE and BM conceived and planned experiments. BM and JH carried out experiments and analyzed data. LL performed immunofluorescence microscopy and analyzed data. BM, JH and SJE prepared figures and wrote manuscript.

## FUNDING INFORMATION

Research was funded by Karolinska Institutet SFO Molecular Biosciences, Vetenskapsrådet (2015-04815), H2020 ERC-2016-StG (715024 RAPID), Ming Wai Lau Center for Reparative Medicine, Ragnar Söderbergs Stiftelse.

## SUPPORTING INFORMATION

**Supplementary Figure 1.**
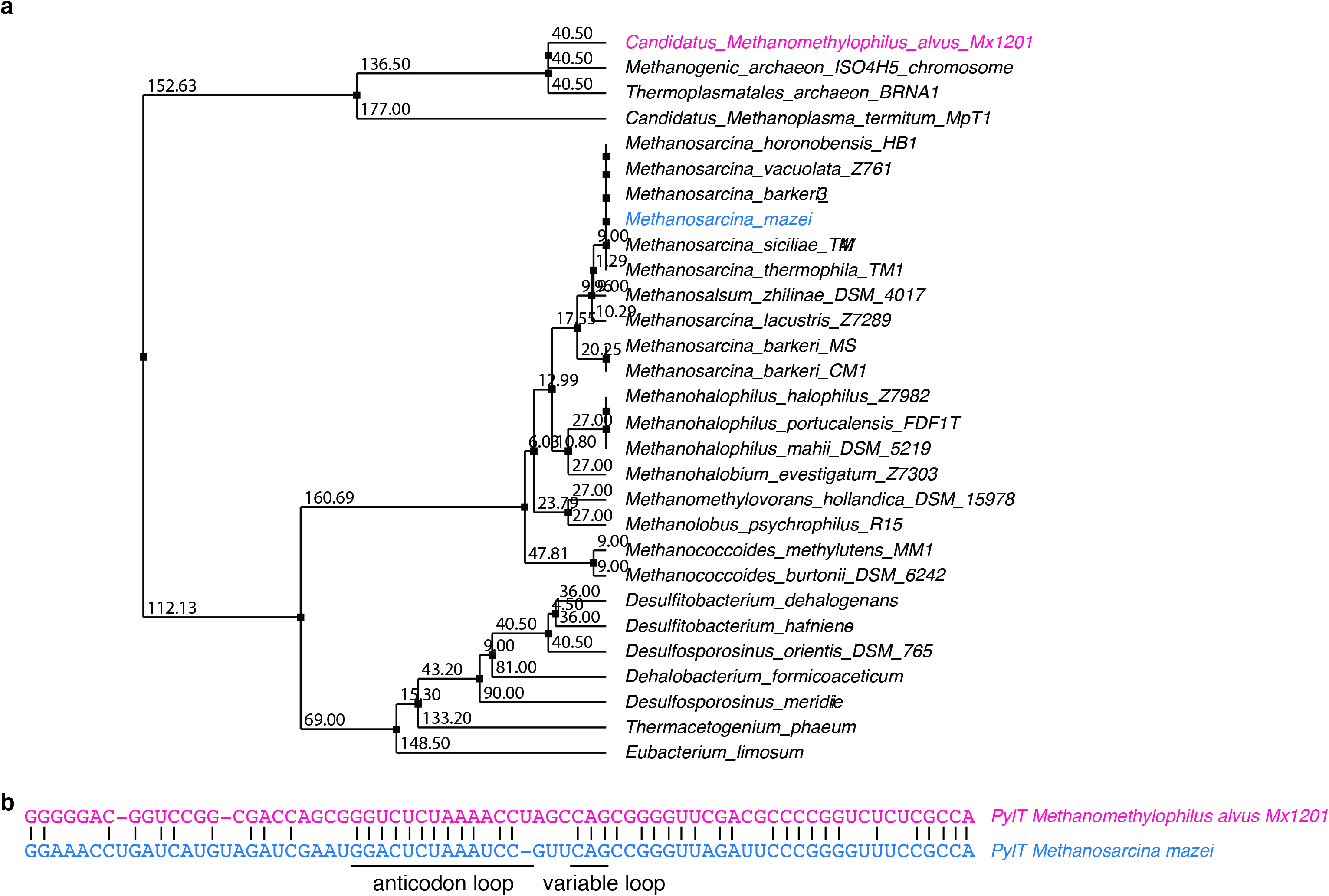
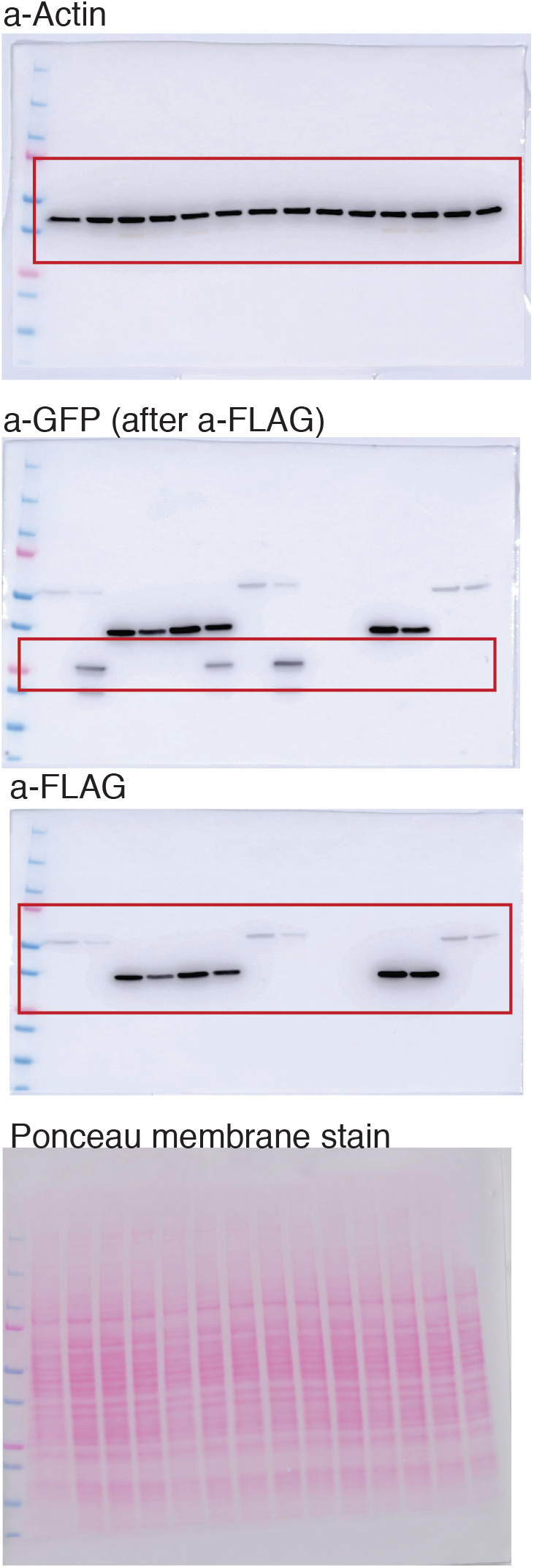
Evolutionary divergence of *Mx1201* PylT. (a) Average distance tree based on a multiple sequence alignment (Tcoffee) of representative PylT sequences of archaeal and bacterial origin. (b) Sequence alignment between *Mx1201* PylT and *Mma* PylT.

**Supplementary Figure 2.**
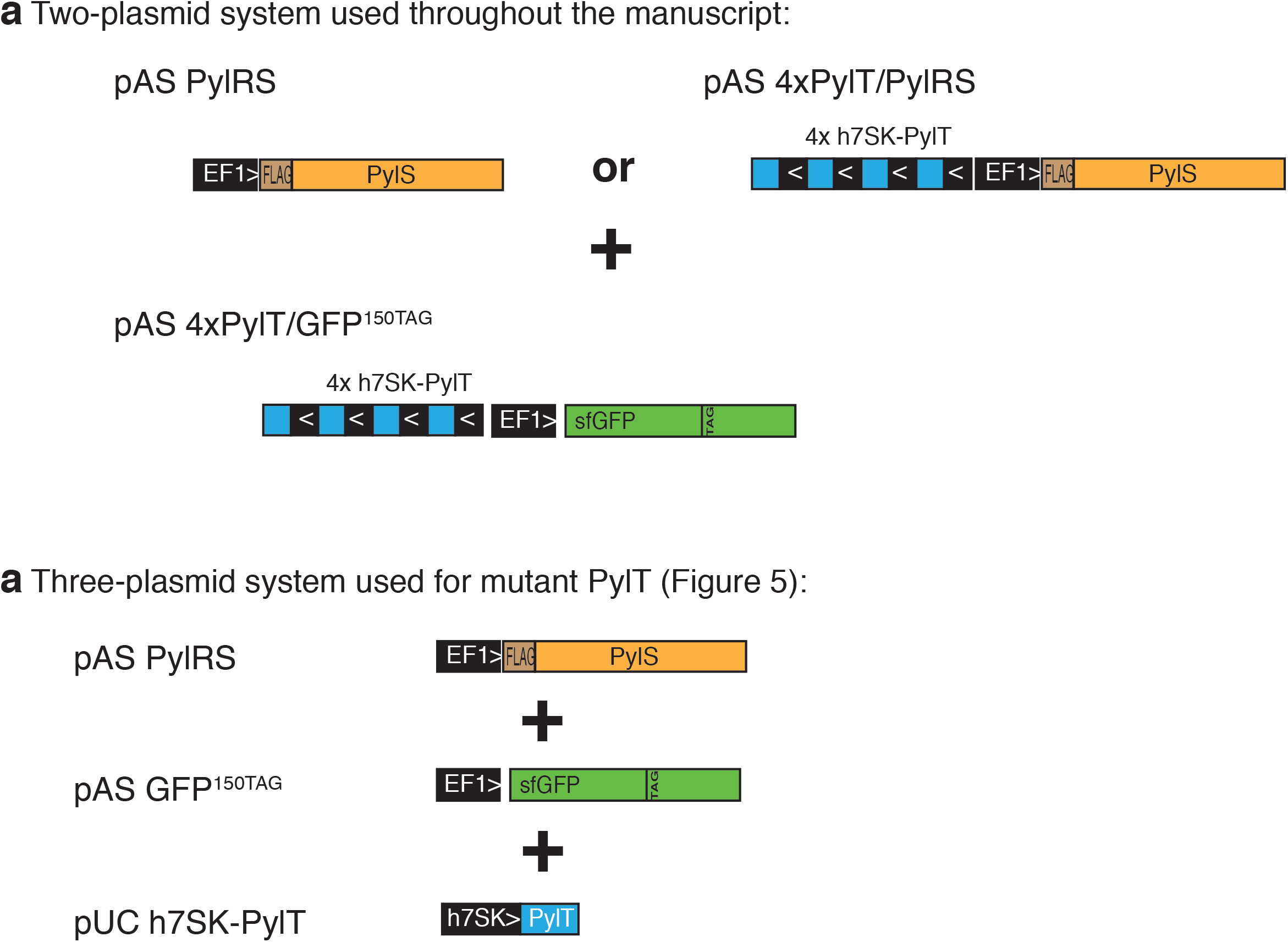
Schematic representation of mammalian expression constructs used in this study. (a) The two-plasmid system comprises a plasmid expressing *PylS* driven by EF1 promoter and a separate plasmid expressing the *GFP^150TAG^* reporter driven by EF1 promoter. The reporter plasmid also carries an expression cassette with four copies of *PylT* driven by h7SK (or U6) promoters. The *PylS* vector may also carry a *PylT* array where indicated. (b) The three-plasmid system comprises separate plasmids for *PylS*, *PylT* and *GFP^150TAG^* reporter. The *PylS* and reporter plasmid are analogous to above but without *PylT*. The PylT plasmid comprises a single copy *PylT* driven by h7SK promoter.

**Supplementary Figure 3.**
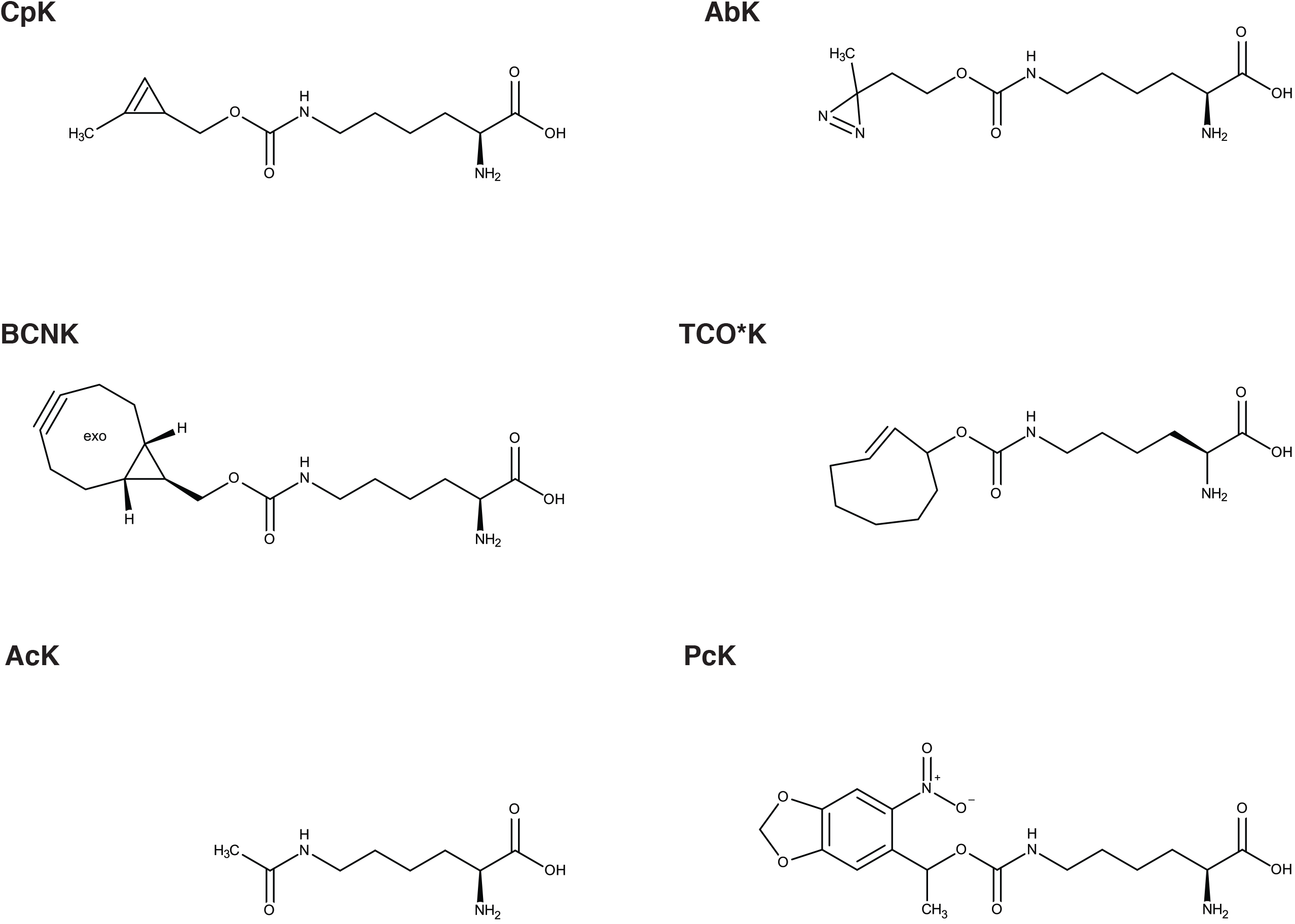
Chemical structures of non-canonical amino acids. Abbreviations used here: Diazirine–L-Lysine (**AbK**); Nε-Acetyl-L-lysine (**AcK**); exo-bicyclo-[6.1.0]-nonyne–L-Lysine (**BCNK**); N6-[[(2-methyl-2-cyclopropene-1-yl) methoxy] carbonyl–L-Lysine (**CpK**); “Photocaged” (2S)-2-(tert-Butoxycarbonylamino)-6-{[1-(6-nitrobenzo[d][1,3]dioxol-5-yl)ethoxy]carbonylamino}hexanoic–L-Lysine (**PcK)**; axial trans-Cyclooct-2-ene–L-Lysine (**TCO^*^K**)

**Supplementary Figure 4.**
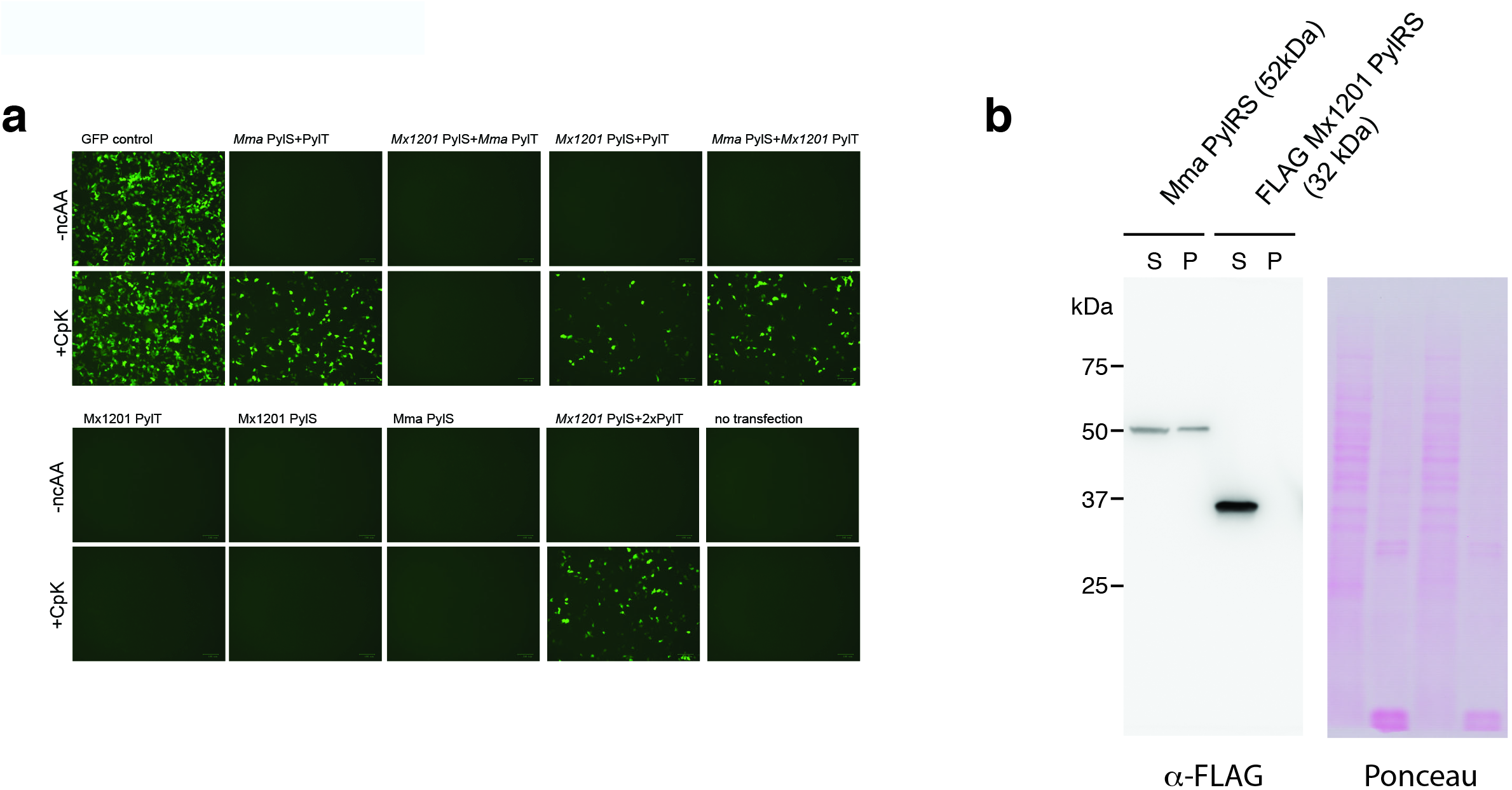
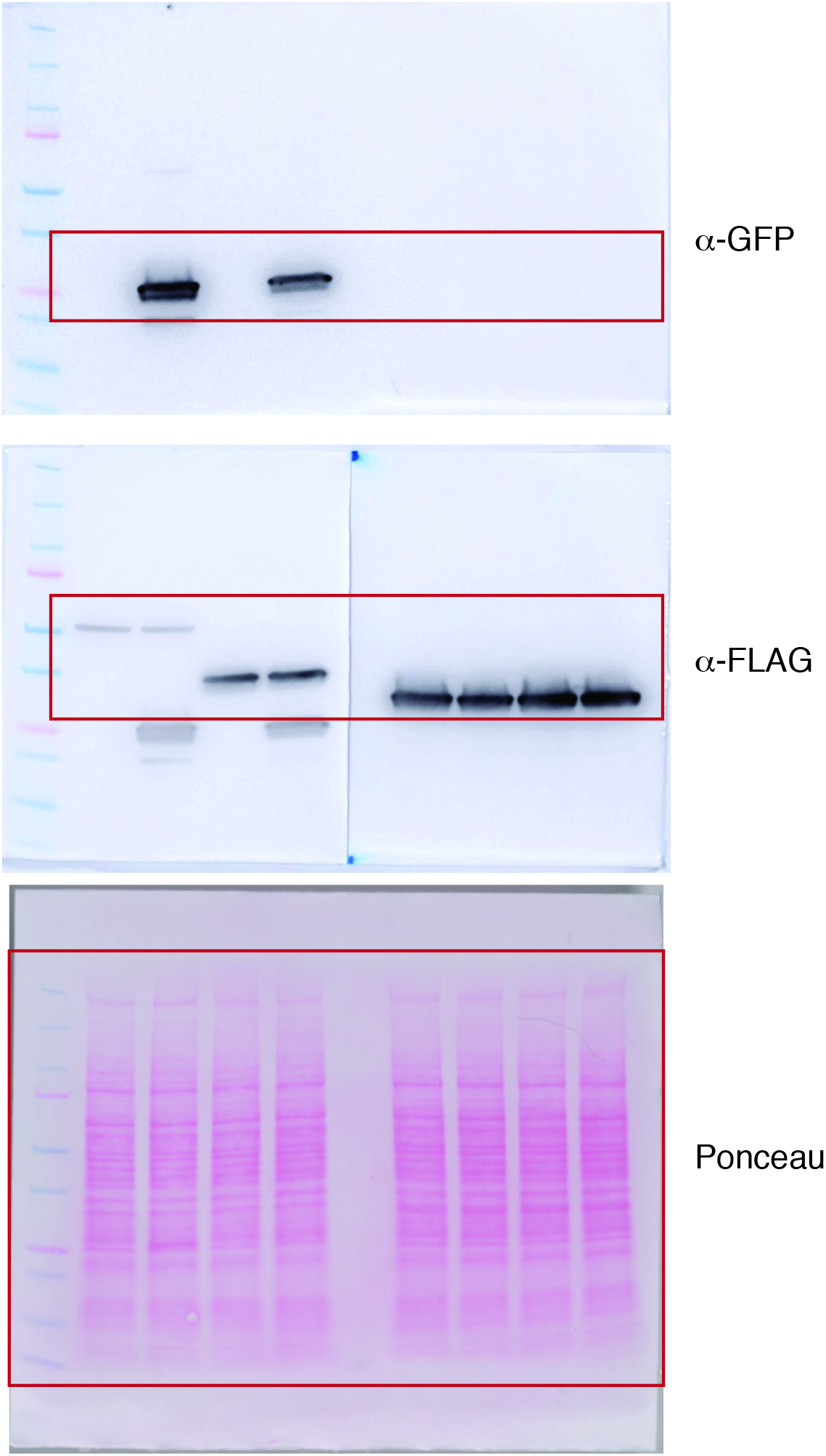
Relating to Figure 2a, b. Fluorescence microscopy images and PylRS solubility test. (a) Fluorescent images of same samples shown in Figure 2a. HEK293T cells were transiently transfected with a GFP^150TAG^ reporter, and a combination of tRNA and synthetase, at a 9:1 ratio. GFP fluorescence is shown as percentage of fluorescence measured with a GFP construct without TAG stop codon (GFPctrl) in the same experiment. For each combination, quadruplicate transfections were performed. For three of the four samples medium was supplemented with 0.2 mM CpK, representative pictures after 24h are shown. (b) Assessment of solubility of *Mma* and *Mx1201* PylRS. HEK293T cells were transfected as in (a). After lysis in RIPA, insoluble material was routinely spun down before western blot. Here the lysate supernatant and insoluble material were brought to the same volume and run side-by-side. Half of the *Mma* PylRS is in the insoluble fraction, whereas *Mx1201* PylRS is soluble.

**Supplementary Figure 5.**
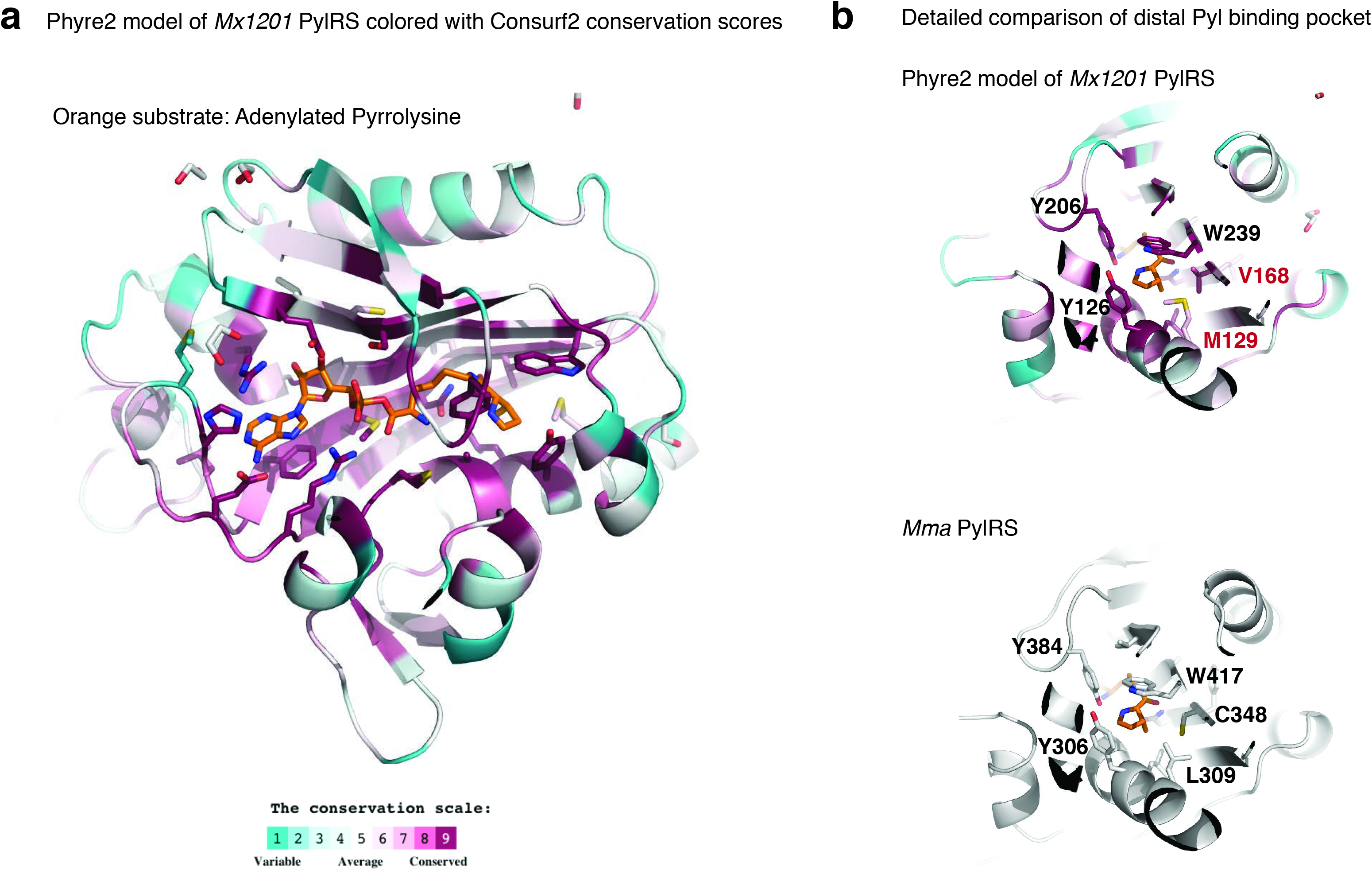
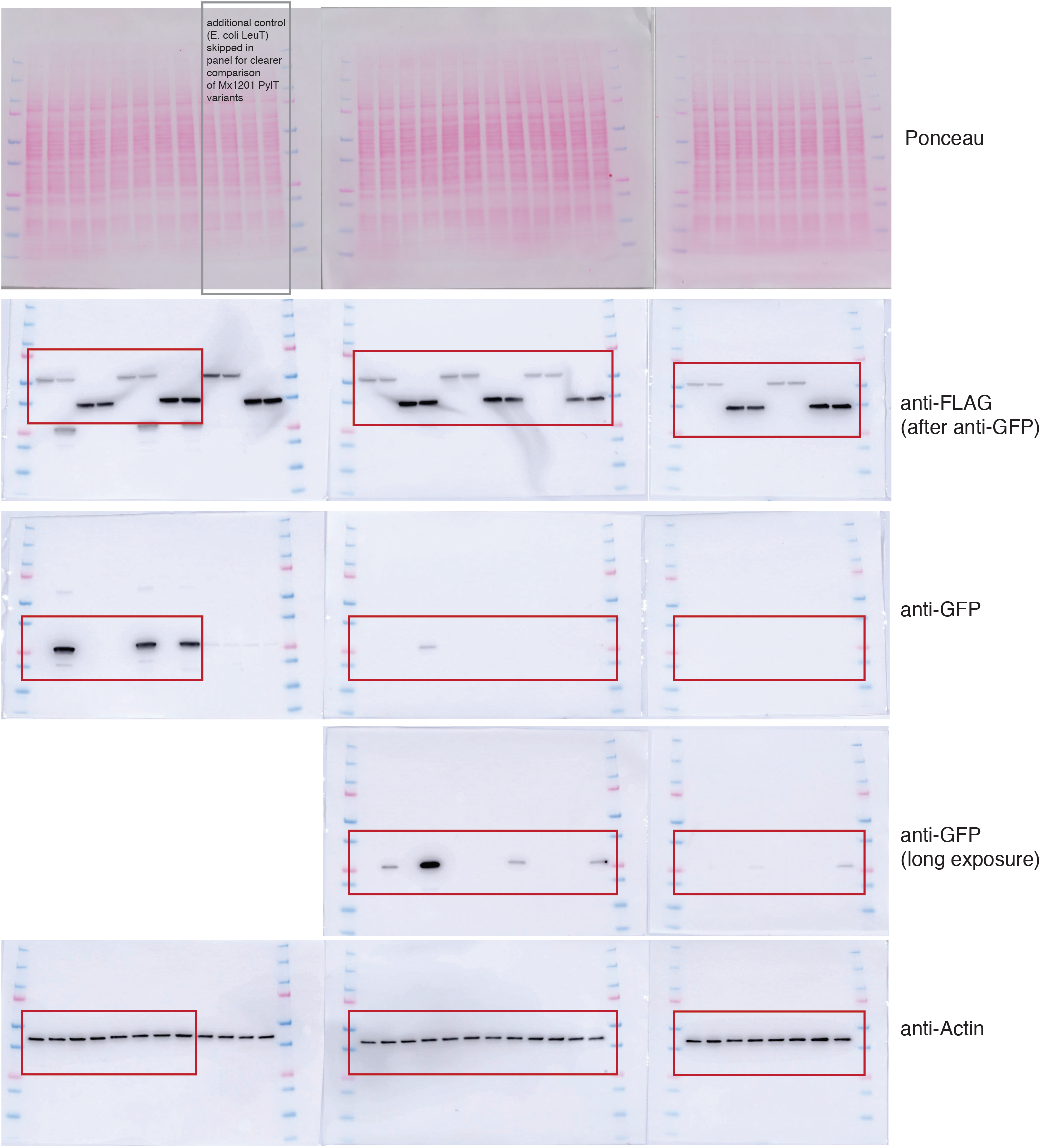
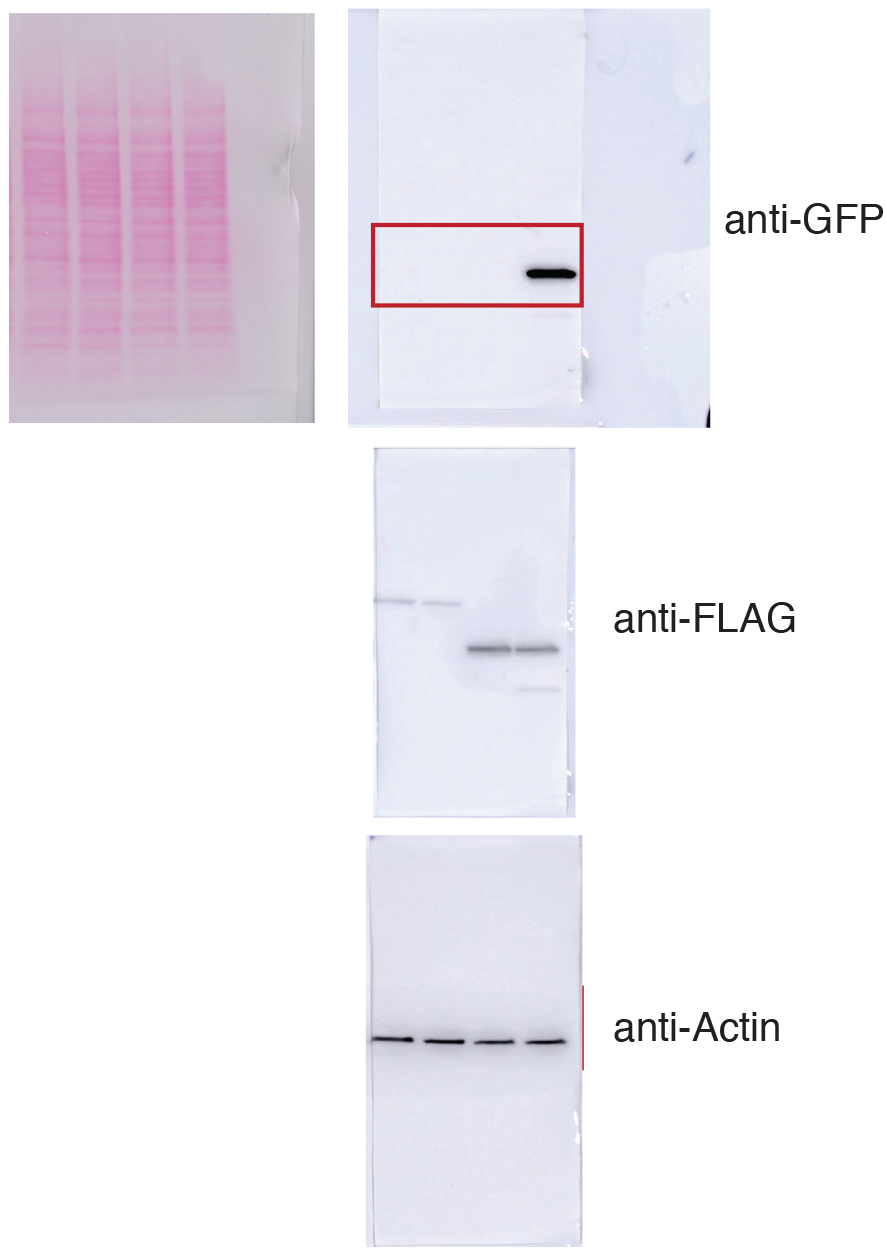
Model of *Mx1201* PylRS protein with Pyl bound to active site. A homology model of *Mx1201* PylRS generated using the Phyre2 web server (http://www.sbg.bio.ic.ac.uk/phyre2/). Per-residue conservation scores were calculated based on multiple sequence alignments using the ConSurf server (http://consurf.tau.ac.il/). (a) Overview *Mx201* PylRS model, colored with conservation scores. Notably, most residues facing the active site are extremely conserved (dark purple), but residues in the outer shells are little conserved. (b) Comparison of Pyl binding by *Mx1201* PylRS and *Mma* PylRS, *Mx1201* PylRS model was aligned to *Mma* PylRS crystal structure complex with adenylated Pyl (PDB 2Q7H). Residues are colored with conservation scores. Red labels indicate *Mx1201*-specific sequence variations in otherwise highly conserved positions

**Supplementary Figure 6.**
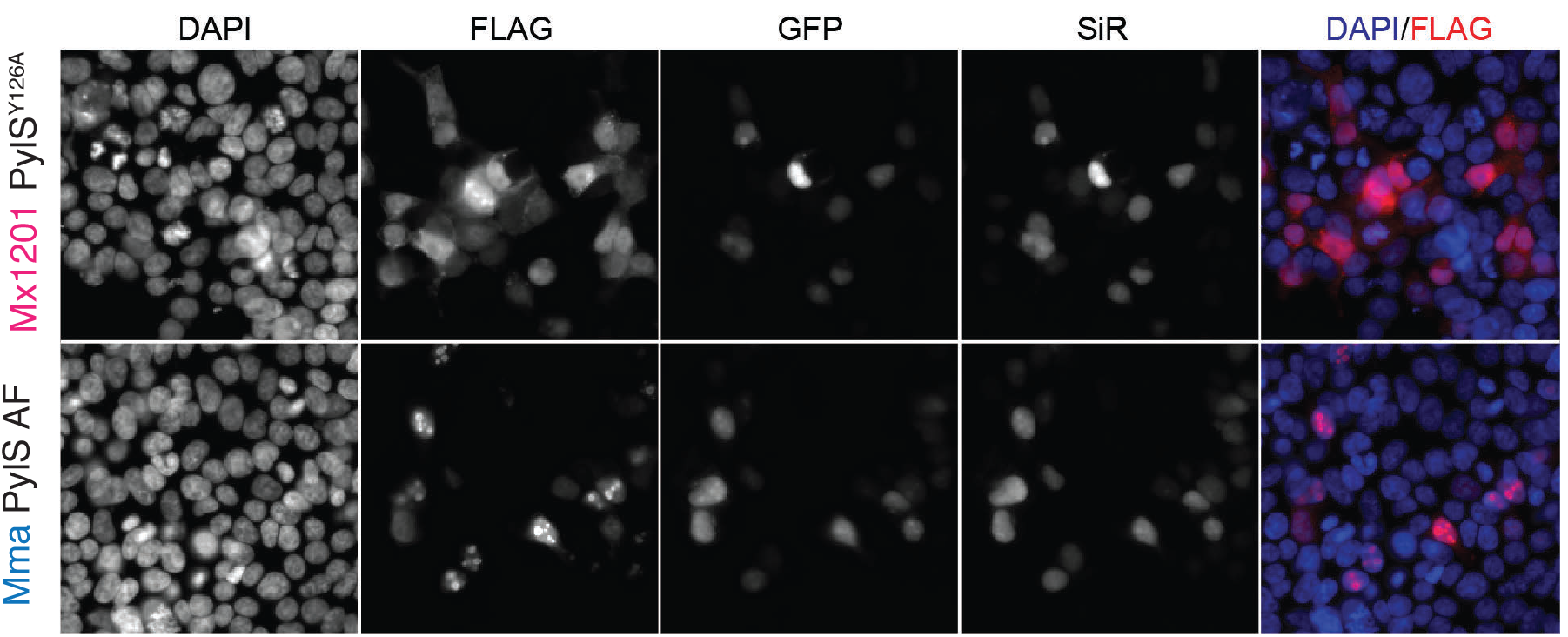
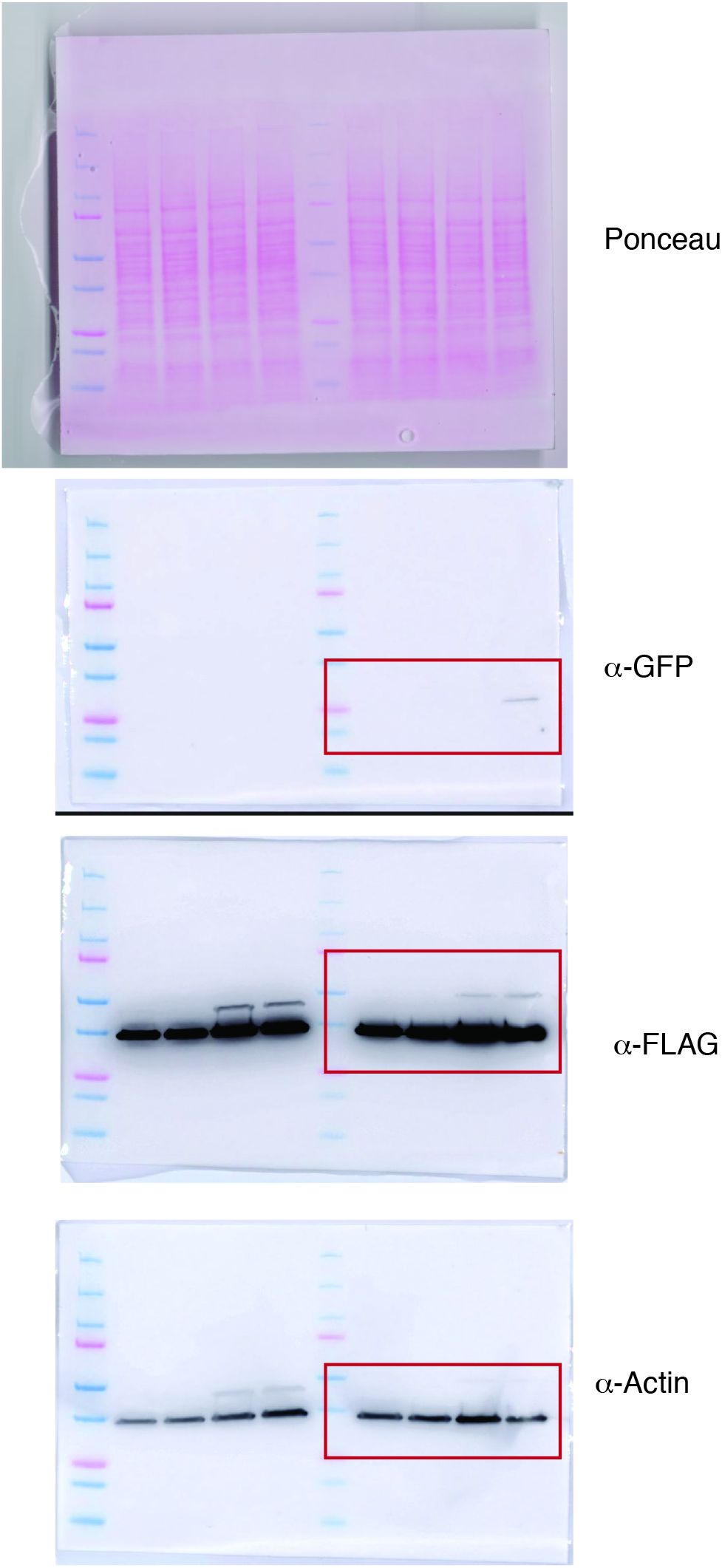
Immunofluorescence microscopy comparing expression of *Mx1201* and *Mma* PylRS. Cells were cotransfected with GFP reporter/4x*Mx1201 PylT*, and *Mx1201* PylRS^Y*126A*^/4x*Mx1201 PylT* and TCO^*^K was added to the growth medium. Cells were fixed and stained using SiR-tetrazine dye. Cells were counterstained with anti-GFP (Alexa 488), anti-FLAG (Alexa 555) and DAPI.

**Supplementary Figure 7.**
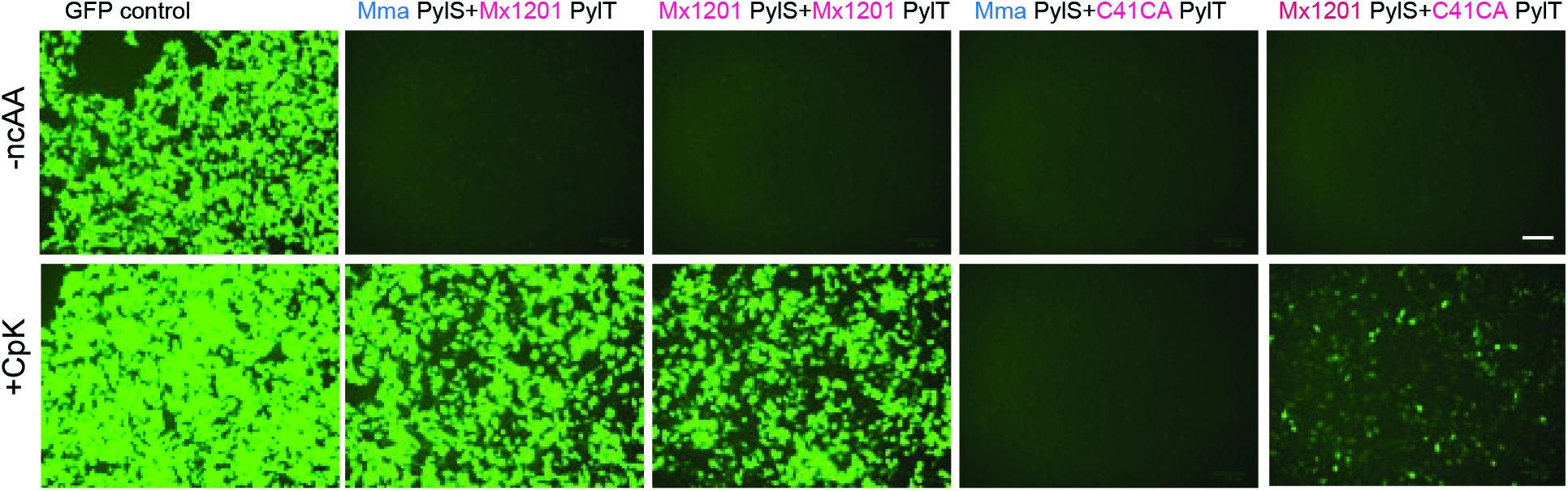
Relating to Figure 5e. Fluorescence microscopy images. Full panel including conditions shown in Figure 5e, showing orthogonality of *Mx1201* PylT^C41CA^ as opposed to wild type *Mx1201* PylT. Also evident is the lower activity with the cognate *Mx1201* PylRS.

## BIBLIOGRAPHY

Ai, H., Shen, W., Sagi, A., Chen, P.R. and Schultz, P.G. 2011. Probing protein-protein interactions with a genetically encoded photo-crosslinking amino acid. Chembiochem 12(12), pp. 1854–1857.

Ambrogelly, A., Gundllapalli, S., Herring, S., Polycarpo, C., Frauer, C. and Söll, D. 2007. Pyrrolysine is not hardwired for cotranslational insertion at UAG codons. Proceedings of the National Academy of Sciences of the United States of America 104(9), pp. 3141–3146.

Baranov, P.V., Atkins, J.F. and Yordanova, M.M. 2015. Augmented genetic decoding: global, local and temporal alterations of decoding processes and codon meaning. Nature Reviews. Genetics 16(9), pp. 517–529.

Borrel, G., Gaci, N., Peyret, P., O’Toole, P.W., Gribaldo, S. and Brugère, J.-F. 2014. Unique characteristics of the pyrrolysine system in the 7th order of methanogens: implications for the evolution of a genetic code expansion cassette. Archaea 2014, p. 374146.

Borrel, G., Harris, H.M.B., Parisot, N., Gaci, N., Tottey, W., Mihajlovski, A., Deane, J., Gribaldo, S., Bardot, O., Peyretaillade, E., Peyret, P., O’Toole, P.W. and Brugère, J.-F. 2013. Genome Sequence of “Candidatus Methanomassiliicoccus intestinalis” Issoire-Mx1, a Third Thermoplasmatales-Related Methanogenic Archaeon from Human Feces. Genome announcements 1(4).

Borrel, G., Harris, H.M.B., Tottey, W., Mihajlovski, A., Parisot, N., Peyretaillade, E., Peyret, P., Gribaldo, S., O’Toole, P.W. and Brugère, J.-F. 2012. Genome sequence of “Candidatus Methanomethylophilus alvus” Mx1201, a methanogenic archaeon from the human gut belonging to a seventh order of methanogens. Journal of Bacteriology 194(24), pp. 6944–6945.

Chatterjee, A., Sun, S.B., Furman, J.L., Xiao, H. and Schultz, P.G. 2013. A versatile platform for single- and multiple-unnatural amino acid mutagenesis in Escherichia coli. Biochemistry 52(10), pp. 1828–1837.

Chatterjee, A., Xiao, H. and Schultz, P.G. 2012. Evolution of multiple, mutually orthogonal prolyl-tRNA synthetase/tRNA pairs for unnatural amino acid mutagenesis in Escherichia coli. Proceedings of the National Academy of Sciences of the United States of America 109(37), pp. 14841–14846.

Chin, J.W. 2017. Expanding and reprogramming the genetic code. Nature 550(7674), pp. 53–60.

Dridi, B., Fardeau, M.-L., Ollivier, B., Raoult, D. and Drancourt, M. 2012. Methanomassiliicoccus luminyensis gen. nov., sp. nov., a methanogenic archaeon isolated from human faeces. International Journal of Systematic and Evolutionary Microbiology 62(Pt 8), pp. 1902–1907.

Elliott, T.S., Townsley, F.M., Bianco, A., Ernst, R.J., Sachdeva, A., Elsässer, S.J., Davis, L., Lang, K., Pisa, R., Greiss, S., Lilley, K.S. and Chin, J.W. 2014. Proteome labeling and protein identification in specific tissues and at specific developmental stages in an animal. Nature Biotechnology 32(5), pp. 465–472.

Elsässer, S.J. 2018. Generation of stable amber suppression cell lines. Methods in Molecular Biology 1728, pp. 237–245.

Elsässer, S.J., Ernst, R.J., Walker, O.S. and Chin, J.W. 2016. Genetic code expansion in stable cell lines enables encoded chromatin modification. Nature Methods 13(2), pp. 158–164.

Gautier, A., Nguyen, D.P., Lusic, H., An, W., Deiters, A. and Chin, J.W. 2010. Genetically encoded photocontrol of protein localization in mammalian cells. Journal of the American Chemical Society 132(12), pp. 4086–4088.

Gunnigle, E., McCay, P., Fuszard, M., Botting, C.H., Abram, F. and O’Flaherty, V. 2013. A functional approach to uncover the low-temperature adaptation strategies of the archaeon Methanosarcina barkeri. Applied and Environmental Microbiology 79(14), pp. 4210–4219.

Herring, S., Ambrogelly, A., Gundllapalli, S., O’Donoghue, P., Polycarpo, C.R. and Söll, D. 2007. The amino-terminal domain of pyrrolysyl-tRNA synthetase is dispensable in vitro but required for in vivo activity. FEBS Letters 581(17), pp. 3197–3203.

Jiang, R. and Krzycki, J.A. 2012. PylSn and the homologous N-terminal domain of pyrrolysyl-tRNA synthetase bind the tRNA that is essential for the genetic encoding of pyrrolysine. The Journal of Biological Chemistry 287(39), pp. 32738–32746.

Kavran, J.M., Gundllapalli, S., O’Donoghue, P., Englert, M., Söll, D. and Steitz, T.A. 2007. Structure of pyrrolysyl-tRNA synthetase, an archaeal enzyme for genetic code innovation. Proceedings of the National Academy of Sciences of the United States of America 104(27), pp. 11268–11273.

Neumann, H., Peak-Chew, S.Y. and Chin, J.W. 2008. Genetically encoding N(epsilon)-acetyllysine in recombinant proteins. Nature Chemical Biology 4(4), pp. 232–234.

Neumann, H., Slusarczyk, A.L. and Chin, J.W. 2010. De novo generation of mutually orthogonal aminoacyl-tRNA synthetase/tRNA pairs. Journal of the American Chemical Society 132(7), pp. 2142–2144.

Nikić, I., Estrada Girona, G., Kang, J.H., Paci, G., Mikhaleva, S., Koehler, C., Shymanska, N.V., Ventura Santos, C., Spitz, D. and Lemke, E.A. 2016. Debugging Eukaryotic Genetic Code Expansion for Site-Specific Click-PAINT Super-Resolution Microscopy. Angewandte Chemie 55(52), pp. 16172–16176.

Nikić, I., Plass, T., Schraidt, O., Szymański, J., Briggs, J.A.G., Schultz, C. and Lemke, E.A. 2014. Minimal tags for rapid dual-color live-cell labeling and super-resolution microscopy. Angewandte Chemie 53(8), pp. 2245–2249.

Nozawa, K., O’Donoghue, P., Gundllapalli, S., Araiso, Y., Ishitani, R., Umehara, T., Söll, D. and Nureki, O. 2009. Pyrrolysyl-tRNA synthetase-tRNA(Pyl) structure reveals the molecular basis of orthogonality. Nature 457(7233), pp. 1163–1167.

Polycarpo, C., Ambrogelly, A., Bérubé, A., Winbush, S.M., McCloskey, J.A., Crain, P.F., Wood, J.L. and Söll, D. 2004. An aminoacyl-tRNA synthetase that specifically activates pyrrolysine. Proceedings of the National Academy of Sciences of the United States of America 101(34), pp. 12450–12454.

Schmied, W.H., Elsässer, S.J., Uttamapinant, C. and Chin, J.W. 2014. Efficient multisite unnatural amino acid incorporation in mammalian cells via optimized pyrrolysyl tRNA synthetase/tRNA expression and engineered eRF1. Journal of the American Chemical Society 136(44), pp. 15577–15583.

Suzuki, T., Miller, C., Guo, L.-T., Ho, J.M.L., Bryson, D.I., Wang, Y.-S., Liu, D.R. and Söll, D. 2017. Crystal structures reveal an elusive functional domain of pyrrolysyl-tRNA synthetase. Nature Chemical Biology 13(12), pp. 1261–1266.

Wan, W., Tharp, J.M. and Liu, W.R. 2014. Pyrrolysyl-tRNA synthetase: an ordinary enzyme but an outstanding genetic code expansion tool. Biochimica et Biophysica Acta 1844(6), pp. 1059–1070.

Willis, J.C.W. and Chin, J.W. 2018. Mutually orthogonal pyrrolysyl-tRNA synthetase/tRNA pairs. Nature Chemistry.

Xiao, H., Chatterjee, A., Choi, S., Bajjuri, K.M., Sinha, S.C. and Schultz, P.G. 2013. Genetic incorporation of multiple unnatural amino acids into proteins in mammalian cells. Angewandte Chemie 52(52), pp. 14080–14083.

Yanagisawa, T., Ishii, R., Fukunaga, R., Kobayashi, T., Sakamoto, K. and Yokoyama, S. 2008. Multistep engineering of pyrrolysyl-tRNA synthetase to genetically encode N(epsilon)-(o-azidobenzyloxycarbonyl) lysine for site-specific protein modification. Chemistry & Biology 15(11), pp. 1187–1197.

